# Accessing pore-blocker bound and alternative conductive conformations of TMEM16A using PIP_2_-assisted adaptive sampling

**DOI:** 10.64898/2026.04.09.717600

**Authors:** Tanadet Pipatpolkai, Ee Hou Yong

## Abstract

Ion channels are promising targets for drug discovery due to their diverse physiological functions. The database of ion channel structures has grown exponentially over the last decade due to advances in structure-determination techniques. However, not all ion channel conformations have been determined, and not all druggable conformations can be modelled as thermodynamically stable under simulation conditions. This greatly limits conformation-specific drug targeting. In this study, we used an endogenous regulator of ion channels, phosphatidylinositol-4,5-bisphosphate (PIP2), as a computational tool to probe the conductive conformation of the TMEM16A calcium-activated chloride channel distinctive from the available structures. By using the PIP2-binding conformation from a coarse-grained model, followed by fluctuation-amplified specific traits (FAST) adaptive sampling in an all-atom configuration, the system transitioned from a closed state to a thermodynamically stable conductive state. The transition also highlights the importance of PIP2 in TM6 helical kink, the opening of the outer gate and the alpha helix on I551. The modelled structure also displays the experimental conductance at sub-300 mV. Using an accelerated weighted histogram (AWH), the binding sites of 1PBC, A9C, niclosamide, and Ani9 pore blockers were determined and validated against previous experimental studies. This paves the way to structure-specific drug development, as overactivation of TMEM16A is correlated with many diseases, such as pulmonary hypertension and ischemic stroke. Together, this study highlights the importance of lipids in stabilising ion channel conformations for targeted drug design and introduces a novel approach to expand the therapeutic targeting of ion channels.

## Introduction

TMEM16A (or Anoctamin-1) is a calcium-activated chloride channel, diversely expressed in the neurones, smooth muscle, and the tracheal epithelial cells ^1^. It is involved in multiple physiological processes, including muscle tone, cell proliferation and epithelial ion transport. Thus, targeting TMEM16A pharmacologically offers a promising avenue for the treatment of multiple diseases, including cystic fibrosis, ischemic stroke, and cancer ^2^.

Multiple synthetic blockers and activators of TMEM16A channel interactions have been characterised using electrophysiological or structural biology approaches. Recent structures of TMEM16A in the Protein Data Bank (PDB) have been determined with activators such as Ca2+ 3,4, gain-of-function mutation (I551A) 5, or with pore blocker 1- hydroxy-3-(trifluoromethyl)pyrido[1,2-a]benzimidazole-4-carbonitrile (1PBC) 6. In addition to the TMEM16A chloride channel, the structure of another protein within the TMEM16 family, TMEM16F, lipid scramblase, has been solved in the apo state and in complex with niclosamide 7. These structures make molecular dynamics (MD) simulation an attractive approach to provide dynamical information on a ns to μs timescale, which has led to multiple recent drug discoveries 8. ^3–8^

TMEM16A activation requires Ca^2+^ ^9–11^ and phosphatidylinositol-4,5-bisphosphate (PIP_2_) as an allosteric modulator ^12–14^. Once Ca^2+^ and PIP_2_ bind to their respective binding sites, they induce a conformational change, a kink at the lower end of TM6 known as the steric gate ^15–17^. The presence of Ca^2+^ also attracts chloride ions, known as the electrostatic gate ^18^. Moreover, Ca^2+^ binding facilitates opening of the outer pore, allowing a TMEM16A-specific drug, such as anthracene-9-carboxylate (A9C), to access its binding site ^19^. Multiple computational studies have proposed an open-state model of TMEM16A with an open outer pore. Those include the model based on TMEM16K scramblase ^19^, the model with PIP_2_ ^20^, and a model which used an electric field to trigger the α-to-π helical transition in TM4 ^21^, which then leads to pore dilation.

In this study, we used PIP_2_, as an endogenous ligand, to drive Fluctuation Amplification of Specific Traits (FAST) adaptive sampling to elucidate the conductive state distinct from the recently solved 1PBC-bound structure of TMEM16A. This work highlights three key features of the conductive state: 1) the opening of the outer gate, 2) the kinking of the TM5 and TM6 helices, and 3) the breaking of the α-helical structure within TM4. The conductance of the model was then calculated to compare to electrophysiological studies. We then used the intermediate and the conductive-state model extracted from the free energy landscape (FEL) to understand the mechanisms of four distinct drugs: 1PBC (1-hydroxy-3-(trifluoromethyl)pyrido[1,2-a]benzimidazole-4- carbonitrile), A9C (anthracene-9-carboxylic acid), niclosamide, and Ani9 (2-(4-chloro-2- methylphenoxy)-N-[(2-methoxyphenyl)methylideneamino]-acetamide), using accelerated weighted histograms (AWH). This approach uncovered previously uncharacterized drug binding sites and accounted for differences in interactions across TMEM16x subtypes. Overall, this work highlights how adaptive and enhanced sampling can recover experimentally elusive, providing structures that serve as novel druggable states beyond the current pool of existing structures.

## Results

### Generating a conductive state of the TMEM16A channel using adaptive sampling

Building on previous studies demonstrating the importance of PIP_2_ as an allosteric activator of the TMEM16A channel ^12–14^, we employed PIP_2_ to facilitate the conformational transition towards the conductive state. To obtain the PIP_2_ binding site, coarse-grained (CG) molecular dynamics simulations were performed using the Ca^2+^ bound structure (PDB entry: 5oyb ^3^) embedded in a POPC bilayer containing 10% PIP_2_ (Supplementary Fig. 1A). The backbone of the CG protein was restrained in x, y, z direction to ensure that the geometry of the cryo-EM structure is maintained. Rather than docking PIP_2_ directly into the known site, this approach locates its binding site through unbiased diffusion within the bilayer. This approach captures the binding pathway and residence behaviour of PIP_2_ rather than assuming a single static pose, which is particularly valuable for a ligand whose highly charged, phosphorylated inositol headgroup makes conventional docking unreliable. The CG simulation was conducted for 10 μs (n=5) and revealed that PIP_2_ binds to the site between TM3, TM4 and TM5, in agreement with multiple previous experimental and computational studies (Supplementary Fig. 1B, C)^14,15,20^. In addition, PIP_2_ remains stably bound upon reaching the binding site (Supplementary Fig. 1D, E). Using coarse-grained free energy perturbation, the binding free energy relative to POPC is at -26.4 ± 0.5 kJ/mol – similar to the PIP_2_ binding site on previously established TREK-1 and Kir channels (Supplementary Figure 2). The convergence of all replicas onto the TM3–TM4–TM5 site, together with the long residence time of the bound lipid (Supplementary Fig. 1D, E), provides support for this site rather than a pre-imposed one. The CG structure was subsequently converted to an all-atoms representation, including PIP_2_ and Ca^2+^, for adaptive sampling (Supplementary Fig. 1F).

We then applied the FAST adaptive sampling technique to promote the transition towards a conductive state by favouring the increase in the distance between the residues lining the porecavity, as detailed in the *Methods* (Fig. 1A). The free energy landscape (FEL) describing the sampled conformation was built using Markov state modelling (MSM), and the frames corresponding to each Perron Cluster Centre Analysis (PCCA) cluster were extracted. Here, the FEL was projected along the time-lagged Independent Component (tIC) based on the tIC Analysis (tICA). By examining the FEL describing the conformational change of the TMEM16A channel opening in the presence of PIP_2_ and Ca^2+^, we observed three clear minima, corresponding to the three states: closed (C), intermediate (I), and conductive (O) (Fig. 1B). We hereafter refer to O as the FAST conductive state, distinct from the 1PBC-bound (7zk3) conformation (see TM3 comparison below). The Mean First Passage Time (MFPT) from C to I, and I to O was found to be 8.9 and 6.4 ms respectively, summed to 15.6 ms, comparable to the previously determined opening kinetics of 20 ms ^22^ (Supplementary Fig. 3A). This is faster than the gating kinetics of the TMEM16A channel in the presence of Ca^2+^ ^22^, thus suggesting that the rate-determining step of the gating event in the TMEM16A channel is controlled by Ca^2+^, with PIP_2_ modulating the extent of channel opening at the outer mouth. Based on the implied timescale analysis and Chapman-Kolmogorov (CK) test, a 10 ns lag time is sufficient to capture all conformational changes required for the transition (Supplementary Fig. 3B, C). Because FAST deliberately reseeds simulations from under-sampled, high-reward states, these regions are not expected to reach full convergence by design. The block analysis also showed that the simulations converging after 15 iterations favors the opening transition, and 10 iterations towards the closing transition (Supplementary Fig. 3D).

**Figure 1.**
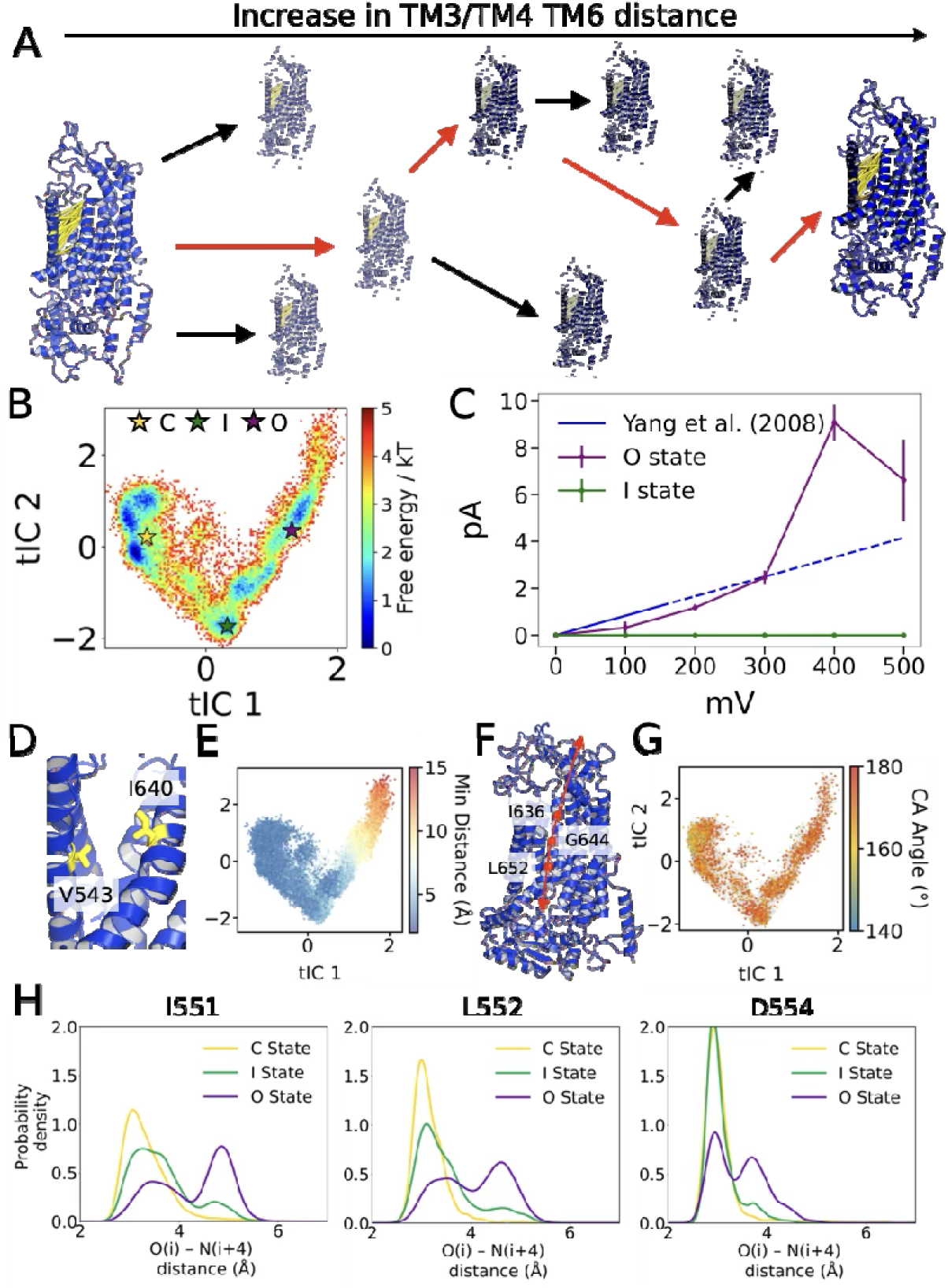
PIP_2_-assisted adaptive sampling led to a conductive state of the TMEM16A. (A) Simplified schematic diagram describing FAST adaptive sampling. The yellow lines represent selected features to increase the distance between TM3/4 and TM6. The red line represents the chosen path. (B) Reweighted free energy landscape corresponding to the TMEM16A conformational transition from closed (yellow star) to intermediate (green star) and open (purple star) states. (C) The calculated current of ion permeation through the channels from 100 mV to 500 mV for the intermediate (green) and open (purple) states. The error bars show the standard deviation (n=3). The blue line represents the current calculated from the experimental conductance value up to 150 mV. Extrapolated values are shown using a dashed line (assuming linear relationship). (D, E) Distributions of distances between V543 and I640 at the outer gate are mapped on the free energy landscape, colored based on their distances. Each dot represents a simulation frame. (F, G) Distributions of the angle on TM6 between I636, G644 and L652 mapped on the free energy landscape, colored based on the angle. Each dot represents a simulation frame. (H) Histograms showing the distribution of i, i+4 distances from each amino acid in I551, L552 and D554 within each PCCA cluster. The C, I, and O states are colored yellow, green, and purple, respectively.

To determine whether the O state is conductive, we assessed the conductance of the TMEM16A channel by applying a positive electric field over a range of +100 mV to +500 mV, in 100 mV increments for 500 ns (n=3). Permeation is infrequent at the lowest potential (100 mV: 0–2 events per replica), so the current there carries a large counting uncertainty; by 200–300 mV the counts rise to 3–9 events and are consistent across the three replicas, indicating a reproducible conductance at these potentials. Per-voltage counting errors are reported in Supplementary Fig. 4A. Ion permeation events were counted and used to calculate the channel current, yielding an I-V relationship similar to the experimental data. At potentials below 300 mV, the O-state conductance closely matches the experimental data within the reported error (Figure. 1C). We then benchmarked this against the most open conformation to date, which is the conformation with 1PBC bound (PDB entry: 7zk3) ^6^. Electric fields at +300 mV and +500 mV were applied across the bilayer and simulated for 500 ns at each membrane potential to calculate the conductance of the 1PBC-bound structure without the drug molecule (n=3) ^23^. The simulations indicate that the channel remains similarly conductive at both +300 and +500 mV (Supplementary Fig. 4B). Notably, a previous study on a similar channel, OSCA, reported that computational studies with electric fields above +400 mV can produce artificially elevated conductance values that substantially exceed experimental expectations ^24^. Overall, the FAST-derived structure reproduces the experimental conductance at sub-300 mV, supporting its assignment as a conductive conformation of the channel. We then explored three TMEM16A conformations with PIP_2_ bound, based on the three PCCA macrostates. A recent MD simulation study highlighted that the recently solved 1PBC-bound structure (PDB entry: 7zk3) is conductive once the pore blocker, 1PBC, is removed from the binding site ^25^. The simulation suggested that the ion flow is inwardly rectifying, in agreement with our work (Supplementary Figure 4). To address similarities between our models, we compare the populations from our PCCA clusters with the simulation of the 7zk3 structure with 1PBC bound. Here, we show that our model did not capture the TM3 axial rotation and the rotation of Y514 away from the pore axis (Supplementary Figure 5A, B). However, our conductive (O) and intermediate-state structures show a partial shift of R515 (Supplementary Figure 5C). When compare the root mean square deviation of each conformation to the structure, the difference are mainly located at the ion conduction pathway, at TM3,4 and 6 (Supplementary Figure 6). This suggested that our model may capture another conductive state, possibly a conformation in which the current-voltage curve is rectifying before becoming fully Ohmic.^25^

The dilation of the outer gate, measured by the minimum distance between the V543 and I640 side chains, is consistent with the similar open-state model for the TMEM16F channel and the model obtained by Jia and Chen (Fig. 1D,E) ^19,20^, with some populations in the I and the O state dilated by 10 and 15 Å, respectively. This emphasises that the simulations sampled the opening of the outer mouth along two tICs. The opening of the outer mouth is accompanied by further bending of TM6 at G644, with a kink angle predominantly at approximately 150°, suggesting that TM6 kinking is associated with channel opening (Fig. 1F,G). Lastly, we examined the possibility of an α-to-π helix transition at the residue on TM5 contributing to channel opening, similar to the mechanism recently reported by Kostritskii et al. 21. (Supplementary Fig. 7). Although we observed the breaking of the α helix in the I and O states at I551 and L552, the formation of the π-helix within the population was not observed. It is also important to note that the α helix at D554 is only broken in the O state (Fig. 1H). This agrees with their mutagenesis data, suggesting the importance of the breakage of the α-helix in TM4 during channel opening 21. Overall, this new model highlighted three key features; the opening of the outer gate, the kinking of the TM6 and the breakage of the α-helix in TM4 is highly associated with TMEM16A opening.^19–21^

### Channel opening is dependent on both PIP_2_ and Ca^2+^

We explored whether the opening of the outer pore, the kinking of TM5 and TM6, and the breakage of the α helix could occur in the absence of PIP_2_ or Ca^2+^. FAST adaptive sampling was conducted using the same starting structure, but without Ca^2+^ (No Ca^2+^), or without PIP_2_ (No PIP_2_), and the corresponding FELs were calculated. Each free-energy landscape was projected onto its own set of time-lagged independent components, since the slowest motions of the channel differ between the PIP_2_- and Ca^2+^-bound conditions. The tIC coordinates, therefore, represent different collective motions in each panel, and the conformational states were identified from the structures populating each minimum. In the absence of PIP_2_, the channel populated only the closed state, with the outer gate (V543–I640) remaining constricted. We also observed a comparable narrowing of the outer gate in the absence of Ca^2+^, indicating that both cofactors are needed to stabilise the conductive conformation. The I and O states were then projected onto the FELs to assess whether they remained accessible under these conditions, confirming that both states were indeed sampled (Fig. 2A). Both No PIP_2_ and No Ca^2+^ simulations converged within the same timescale as the simulations with PIP_2_, with a lag time of 10 ns and within the same number of iterations (Supplementary Fig. 8). When Ca^2+^ was absent, the minimum corresponded to the closed state, and the I and O states were located in the high FE region, as expected (Fig. 2A). Interestingly, the simulation with Ca^2+^ shows an I-state structure at the minima, suggesting that Ca^2+^ assisted the opening of the outer gate to some extent. It is also clear that, in the absence of Ca^2+^, the conductive state is inaccessible. Even if they are accessible, they are located in a region of higher free energy, suggesting that they are unstable. Overall, these FELs highlighted the crucial role of PIP_2_ in stabilising the conductive state of TMEM16A and reducing the energy barrier for the conductive-state transition.

**Figure 2.**
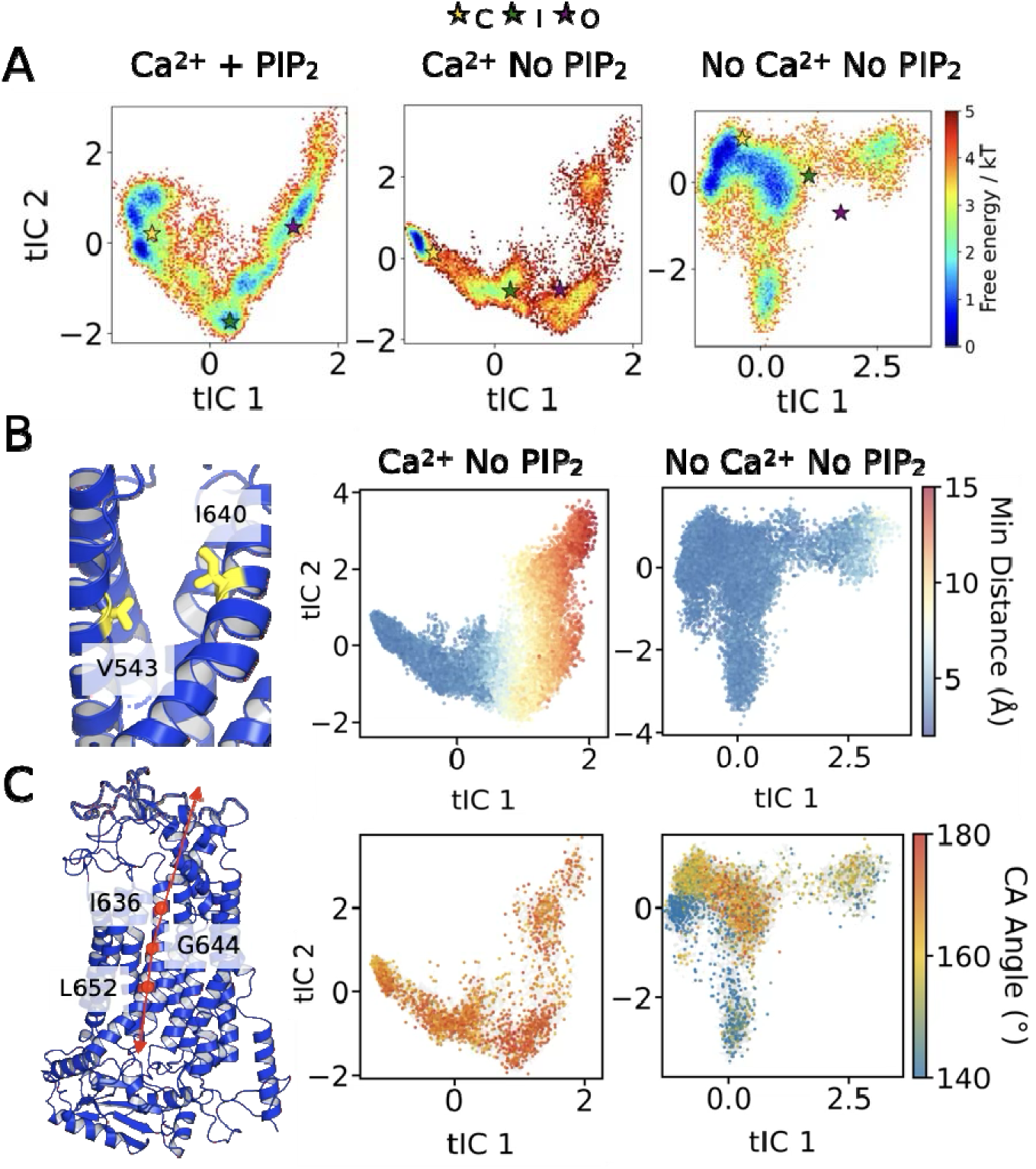
Free energy landscape of pore opening in the presence and absence of PIP_2_ or Ca^2+^. (A) Reweighted free energy landscape corresponding to the TMEM16A conformational transition from closed (yellow star) to intermediate (green star) and open (purple star) states. Landscapes are shown in the presence of PIP2 and Ca2+ (Left), with Ca2+but no PIP2 (Middle), or without any ligands (Right). (B) Distributions of distances between V543 and I640 at the outer gate, or (C) the angle on TM6 between I636, G644, and L652, are mapped on the free energy landscape Ca2+but no of PIP2 (Middle) or without any ligands (Right). Each dot is colored based on its distance or angle, respectively, representing a frame from the simulations.

Next, we sought to investigate the features of the O state in the FEL in the absence of Ca^2+^ or PIP_2_. The two key features, namely, outer gate distance and TM6 helical bending, were calculated based on the structure without Ca^2+^ or PIP_2_ (Fig. 2B,C). Even when the opening was plausible, the conductive state did not correspond to the minimum (Fig. 2A, B). In the simulation with Ca^2+^, the opening towards the I state was plausible, but not towards the O state, as the structure was not located at a minimum. However, the simulations explored an alternative conductive state. All these plausible states are located in regions of the free energy landscape with high free energy, suggesting that the structure is highly unstable. Even more so in simulations with no Ca^2+^, the open-outer-gate structure was not observed at all during sampling. This suggests that Ca^2+^ is crucial for opening and accessibility of the outer mouth, and that PIP_2_ lowers the depth of the energy well in the open conformation.

The second parameter of consideration is the helical bending at TM6 (Fig. 2C). The results highlighted that both simulations with and without PIP_2_ sampled the same helical bending conformational space (between 160° to 180°). This was not observed in the absence of Ca^2+^, where the conformation is mostly sampled below 160° (Supplementary Figure 9). Interestingly, there was a bending at a region with TM6 bending at 160°, yet no outer-gate opening (tIC1 ∼ 0, tIC2 ∼ 0). This observation suggests that the opening of the outer gate and the helical bending may not be tightly coupled processes.

Overall, using PIP_2_ assisted the FAST sampling process in reaching the conductive state by favouring the energy minima at that state. This method provides a powerful computational tool for accessing conformations that have not yet been experimentally resolved.

### 1PBC and A9C bind to the intermediate-open conformation of TMEM16A

At this point, we have successfully captured the conductive state of TMEM16A using PIP_2_-assisted adaptive sampling. To further investigate the conductive state, we studied its pharmacology using the TMEM16A-specific pore blockers 1PBC (Fig. 3A) and A9C (Fig. 3F). Accordingly, we aligned the O-state model with the TMEM16A 1PBC-bound structure (PDB entry: 7zk3) and transposed the 1PBC molecule onto the O state. Unexpectedly, 1PBC dissociated from the binding site in all three replicas, with the ligand RMSD rising above the 6 Å dissociation cutoff in each case (Fig. 3B). This contradicts previous findings and raises the question of whether this open state represents a non-physiological state of the channel. We hypothesised that the inhibitor binding involves an “induced-fit” conformational change of the channel and therefore redirected our analysis to the intermediate state. Accelerated weighted histogram (AWH) was used to enhance the sampling of the 1PBC molecule along the z-axis within the ion permeation vestibule (Fig. 3C). This approach has been used to determine the binding site of astemizole on the hERG channel in a previous study, and similarly with well-tempered metadynamics to determine the binding site of lidocaine on the Nav1.4 channel^26,27^. Using AWH, we found that the binding site is located 15 Å above K645 (Fig. 3D). The I state is non-conductive (Figure 1C). The barrier height is very shallow to the bulk, likely due to the lack of rotation on the TM3. Despite the lack of convergence in the simulations, a rough estimate of the drug-binding site location can be obtained using this methodology (Supplementary Fig. 10A). Using ProLif, the residues that interact with 1PBC were analysed, and results showed that 1PBC predominantly forms Van der Waals interactions with R535, T539 and M518 (Fig. 3E, Supplementary Figure 11). These residues are not exactly the same as the ones identified in the structural studies. Despite differences in our binding mode, our results partially agree with previous mutagenesis data, as mutations at these three residues significantly affect the IC_50_ of 1PBC. ^6^

**Figure 3.**
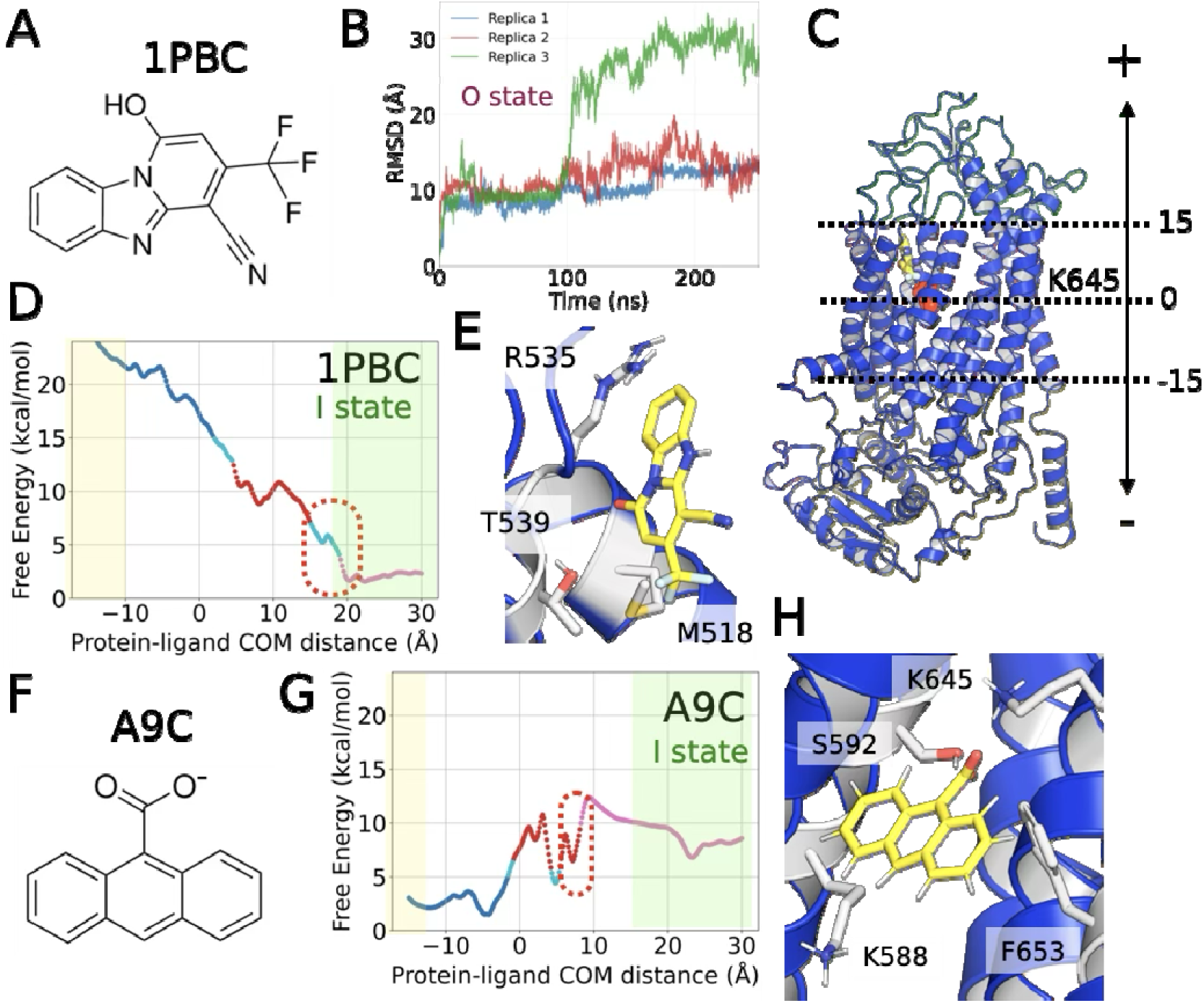
1PBC and A9C bind to the intermediate state of TMEM16A. (A) Chemical structure of 1PBC. (B) RMSD calculation of 1PBC in the conductive state TMEM16A channel, fitted to the Cα atoms. Different colors indicate different repeats from the simulation. (C) Schematic representation of the AWH simulation on the TMEM16A (blue cartoon) channel using the z-axis distance between the center of mass of K645 (spheres) and the center of mass of the drug molecule (yellow sticks). The positive CV (green) indicates the extracellular region, whereas the negative CV (yellow) indicates the intracellular region. (D, G) Average free energy landscape extracted from the AWH simulations across 1.30, 1.35, 1.40, 1.45, and 1.50 μs simulations for the sampling of (D) 1PBC or (G) A9C along the z-axis of the pore. The simulations were conducted in 4 windows with bias shared among them. Blue, red, and pink represent the clusters extracted using DBSCAN with □ = 0.25 and a minimum cluster size of 15. (E, H) Representative structures of the population from the red cluster, with contacting residues shown in white. The 1PBC or A9C is shown in yellow. (F) Chemical structure of A9C.

Next, we applied the same approach to test a second, small-pore blocker, A9C (Fig. 3F). Given that no structure of the drug-bound conformation is available, AWH was used to sample the A9C-bound conformation. The binding site is in a distinct minimum with an energy barrier of ∼5 kcal/mol. From the energy minimum, ProLif exhibits clear minima at a site near the central cavity, with K645 in agreement with the previous electrophysiological study (Fig. 3G,H, Supplementary Figure 11) ^19^. The binding site is below the site obtained from the TMEM16K-derived model. In this site, the existence of the minimum seems to be better converged than that of 1PBC (Supplementary Fig. 10B). This explains why the binding of the smaller ligand fits with our induced-fit model and supports the hypothesis that the pore-blocker-bound conformation is the intermediate state, whereas the fully conductive state exists in a different conformation^19^.

### Inhibitory mechanism of niclosamide

There is an ongoing debate about whether niclosamide is an activator or an inhibitor of the TMEM16A channel ^28–30^. In this work, the inhibitory mechanism of niclosamide is the main focus. First, we made the assumption that the inhibitory binding site of niclosamide on TMEM16A is the same as that of TMEM16F ^7^. Thus, the coordinates of niclosamide from TMEM16F were transferred to both the intermediate and conductive-state models of TMEM16A, and equilibrium MD simulation was conducted for 250 ns (n=5). The RMSD of the drug within the binding pocket was calculated, and it is clear that niclosamide does not stably reside in the binding pocket, as evidenced by the high RMSD (Fig. 4A). Additionally, in the conductive state, simulations at +500 mV show multiple ion permeation events across the bilayer, inconsistent with niclosamide acting as an inhibitor of TMEM16A (Fig. 4B).

**Figure 4.**
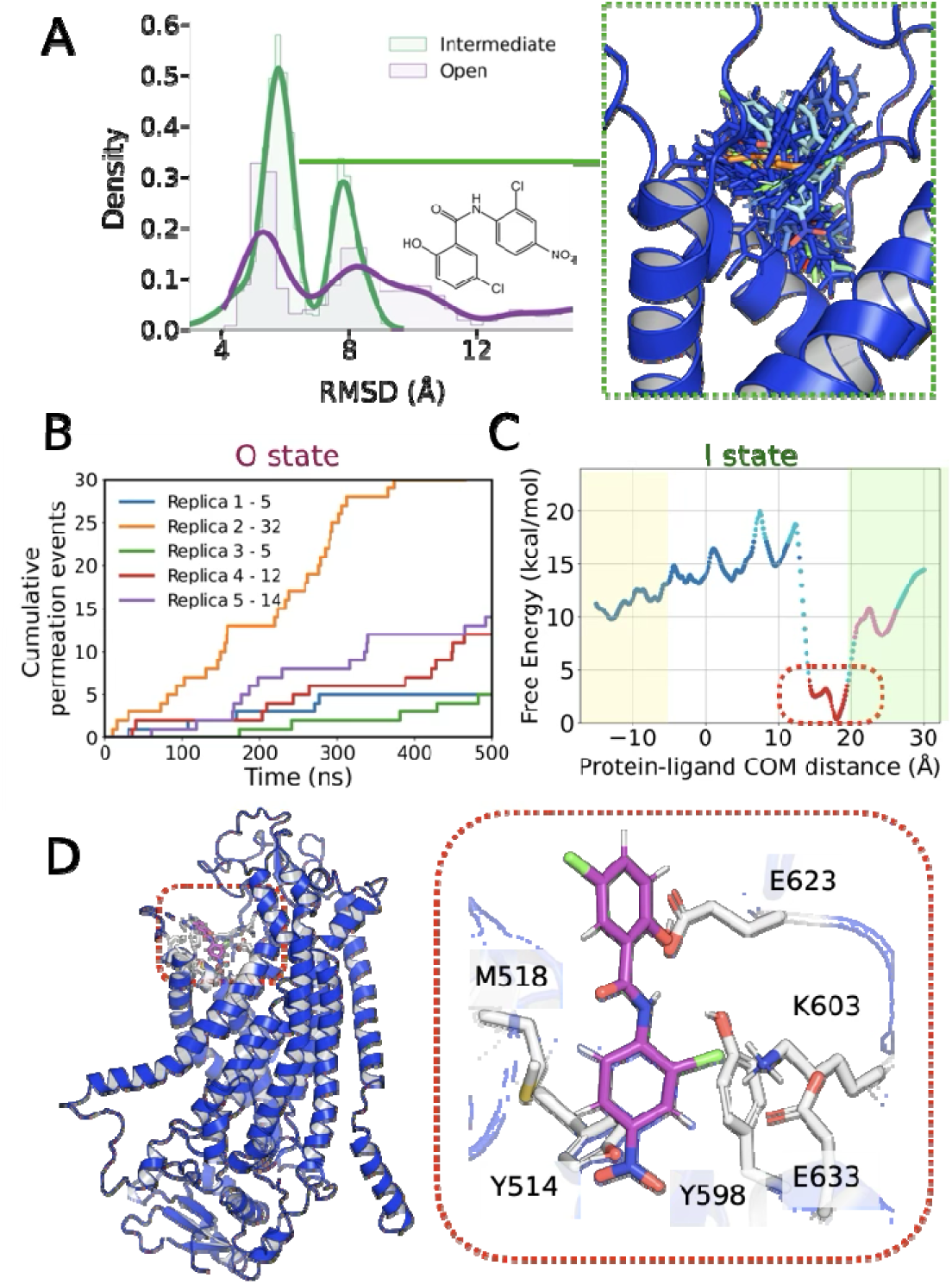
Niclosamide binds at the outer gate of TMEM16A. (A) (Left) Histogram describing the RMSD calculation of niclosamide in the conductive state (purple) and the intermediate state (green) of the TMEM16A channel, fitted to the C_α_ atoms (n=5, 250 ns each). (Right) Different colours indicate different snapshots from the frame cluster from 5 repeats of 250 ns of the simulations. The grey surface represents the structure from the minima derived from AWH simulations. (B) Cumulative permeation of chloride ions across the O state TMEM16A with niclosamide bound under a +500 mV electric field. Different colours indicate different repeats. (C) Average free energy landscape extracted from the AWH simulations across 1.30, 1.35, 1.40, 1.45, and 1.50 μs simulations for the sampling of niclosamide along the z-axis of the pore. The simulations were conducted in 4 windows with bias shared among them. Blue, red, and pink represent the clusters extracted using DBSCAN clustering with □ = 0.25 and minimum population per cluster = 15. (D) Representative structures of the population from the red cluster, with contacting residues shown in white. Niclosamide is shown in purple.

Previous studies suggested that inhibition of the TMEM16A channel by niclosamide is independent of PIP_2_, suggesting that the channel is likely in an intermediate state, rather than a fully open state ^28^. We then investigated the effect of niclosamide binding when the TMEM16A channel is in an intermediate state. From the AWH FEL of I state TMEM16A with niclosamide, we observed a clear free-energy minimum at the outer mouth of the pore, suggesting a well-defined binding site (Fig. 4C). At this binding site, ProLif analyses show Van-der-Walls interaction between M518, E633, K603, Y514, Y598 and E623 (Fig. 4D, Supplementary Fig. 11). The second nitrophenyl ring is supported by K603 on the chlorine atoms, while Y514, and E633 interact with the nitro group, providing stability. Notably, this is a different site from the niclosamide binding site of the TMEM16F scramblase (Supplementary Fig. 12A). To test the stability of niclosamide in the TMEM16F binding pose, we conducted 500 ns simulations of the TMEM16F construct with niclosamide bound. Here, our simulations show that the site obtained from cryo-EM dissociates with RMSD > 15 Å (Supplementary Figure 12B). Thus, this suggests that the binding is unstable, and it is possible that the niclosamide binding site will be different in TMEM16A. Interestingly, the niclosamide binding site on TMEM16A (i.e., Y514, M518 (T518 in mouse), K603 and E633) is highly conserved. (Supplementary Fig. 12C). This aligns well with previous mutagenesis data, which suggested that mutations at T606, T607, and T610 in TMEM16F reduce its scramblase activity, given their close proximity to E633 in the TMEM16A binding site. This binding pose is also different from the reported docked pose, which has been shown to be activatory^29^. Together, these highlight how the same drug can act via different modes of channel inhibition (TMEM16A vs TMEM16F) and how conserved residues might be involved in the same inhibitory function across subtypes ^7,28,30^.

### Ani9 specifically targets TMEM16A but not TMEM16B at the outer gate

Ani9 is a specific TMEM16A chloride channel blocker that lacks blocking activity on TMEM16B (Supplementary Fig. 13A). Inspired by the previous dual mechanism of Tetraethylammonium (TEA) blocking, which has inhibitory action from both the intracellular and extracellular sides ^31^, we set up two hypotheses: (1) Ani9 binds and blocks the outer gate of the channel when the outer gate of the channel is dilated by PIP_2_; hence why TMEM16B, which has no outer gate, is not affected by the drug. (2) Ani9 has an intracellular binding site, which is not conserved between TMEM16A and TMEM16B.

First, the plausibility of Ani9 as an extracellular blocker was tested by placing the drug in the outer pore cavity of the conductive state TMEM16A and conducting equilibrium MD simulations under a +300 mV electric field for 500 ns (n=3). Here, the simulation showed that the drug stably resides in the binding pocket with relatively low RMSD. (Fig. 5A). In addition, Cl- permeation through the pore was not observed, suggesting that this binding mode is plausible as a pore blocker in the conductive state of the channel (Fig. 5B). On this site, ProLif analyses show that. The amide bond in Ani9 interacts with Q637 and Y514, and the drug forms a Van-der-Walls interaction with M518, K603 and E633 (Fig. 5B,C). However, sequence alignment between TMEM16A and TMEM16B displayed no vital difference in the sequence at the outer gate of the channel, particularly between residues 541 and 660, but showed a minor difference in the lower gate of the channel (Supplementary Fig. 13B, C). Thus, we propose that the inner gate in TMEM16B is not wide enough for the drug to bind, but is wide enough for the channel to conduct. This is further supported by the channel’s much lower unitary conductance of 0.8 pS. To validate this point, FAST adaptive sampling was applied on the conformation with Ca^2+^ but without PIP2 to attempt to open the channel using the AlphaFold3 model of TMEM16B. Here, we show that using the same setup, no minima are observed in the distortion of the widening of the outer gate compared to the TMEM16A channel (Fig. 5D, 5E, Supplementary Figure 14). Together, this demonstrates that FAST can be used to assess the plausibility of the druggable state using an AI-generated structure as an initial template.

**Figure 5.**
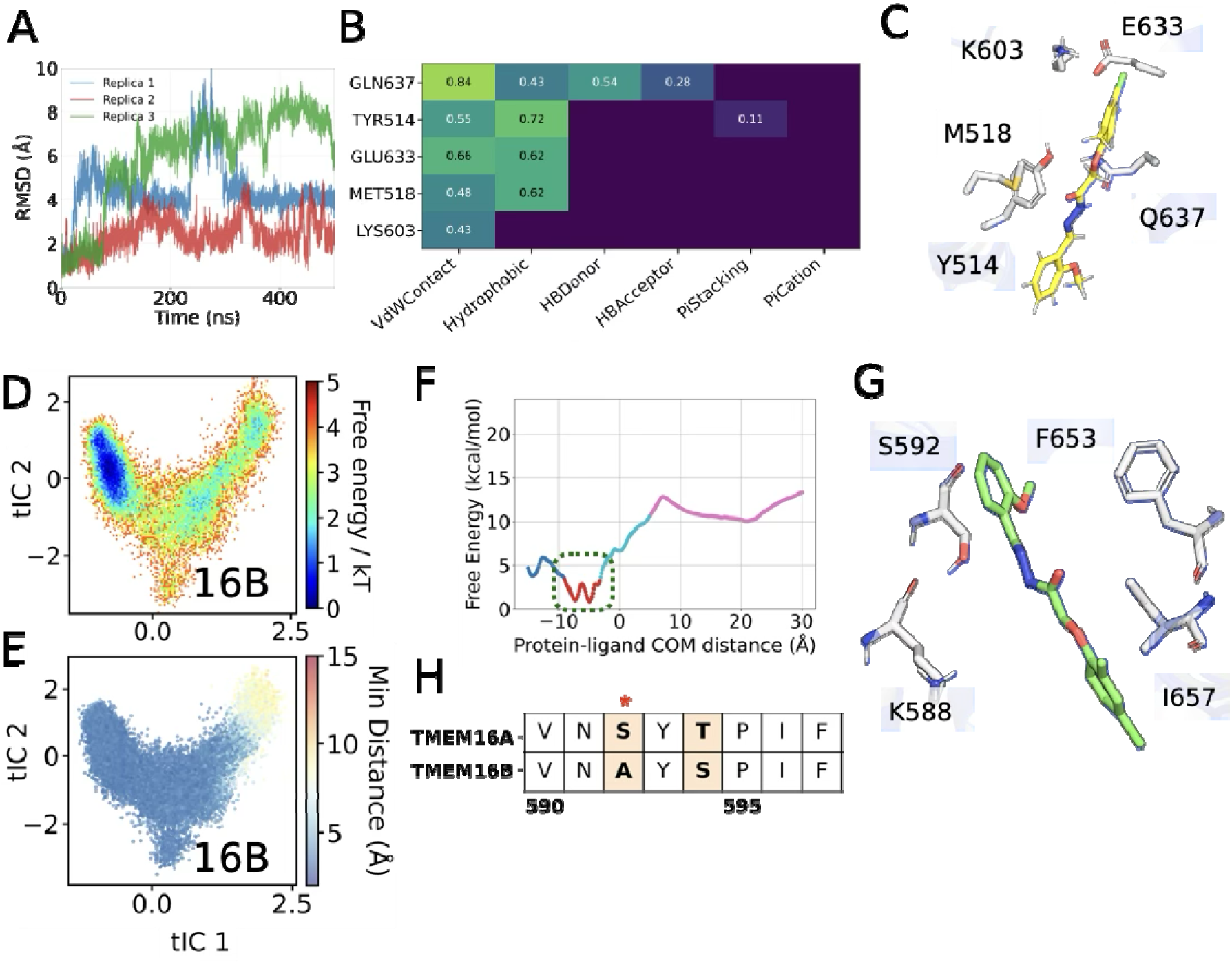
Two plausible binding sites of Ani9 on TMEM16A. (A) RMSD calculation of Ani9 in the conductive state TMEM16A channel, fitted to the Cα atoms. Different colors indicate different repeats from the simulation. (B) ProLIF profile of residual interaction between Ani9 and the protein. (C) Representative structures of the population from the equilibrium simulation where Ani9 (shown in yellow) is placed at the outer mouth of the pore, with contacting residues shown in white. (D) Reweighted free energy landscape corresponding to the TMEM16B conformational transition with Ca2+. (E) Distributions of distances between V543 and I640 at the outer gate (F). (Left) Average free energy landscape extracted from the AWH simulations across 1.30, 1.35, 1.40, 1.45, and 1.50 μs simulations for the sampling of Ani9 along the z-axis of the pore. The simulations were conducted in 4 windows with bias shared among them. Blue, red, and pink represent the clusters extracted using DBSCAN with □ = 0.25 and a minimum population per cluster of 15. (Right) Representative structures of the population from the green cluster, with contacting residues shown in white. Ani9 is shown in green. (H) Sequence alignment between TMEM16A and TMEM16B using the Needleman-Wunsch algorithm, with BLOSUM62 scoring matrix. The orange region indicates a partially conserved region. The asterisk marks S592. (G) Representative structures of the population from the minima of the AWH simulation where Ani9 (shown in green) is located at the intracellular region, with contacting residues shown in white.

As for the possibility of drug binding at the inner gate, the intermediate state TMEM16A structure, which has a closed outer gate, was used. Then, AWH was used to sample the drug conformation. The FEL highlighted that a minimum is located at the inner gate, differentfrom all other tested drugs (Fig. 5F). Using ProLIF analysis, within the binding site, S592 interacts with the amide group via Van-der-Waals interactions, and F653, I657, and K588 form a hydrophobic pocket around the drug molecules (Figure 5G, Supplementary Figure 11D). The only residue which is not conserved between TMEM16A and TMEM16B in this binding site is S592 (Fig. 5G). Thus, the hydrogen bond between S592 and the amide bond might be the crucial interaction conferring specificity between Ani9 and TMEM16A, leading to a second possibility that Ani9 might bind to the channel intracellularly.

## Discussion

In this study, PIP2-assisted FAST adaptive sampling was used to gain insight into a conformation currently inaccessible to experimental structure-determination methods. Previously, FAST has been widely used to probe the cryptic pocket of a drug ^32^, protein folding ^33^, the open state of the GABAA receptor ^34^, and the ADAM10 sheddase mechanism ^35^. This study expands the power of the known endogenous ligand PIP_2_, combining with the adaptive sampling method to obtain the conductive state structures with three key characteristics: 1) the wider outer mouth, 2) the kink at G644 on the TM6, and 3) the widening of the lower gate and the breakage of the α-helix on the TM6 at I551, in agreement with previous experimental and computational studies. However, this study did not capture the α-to-π transition likely due to conformational heterogeneity within the energy minima. This suggests that the formation of the π-helix may not be crucial for opening the channel’s outer gate. This is also observed in the OSCA mechanosensitive chloride channel ^24^. At up to +300 mV, the conductance of the FAST-derived conductive state also matches that of the experimental data. The deviation at higher potentials is likely due to the distortion of the protein structure caused by the electric field, resulting in an overestimation of the unitary conductance, similar to what was observed in TMEM63A ^36^.

We investigated whether the most open conformation of the channel is, in fact, the one in which the blocker is bound. By performing a series of enhanced-sampling simulations and equilibrium MD, our results indicate that the FAST conductive state has a different pore geometry from the drug-bound state, in particular at TM3; however, the intermediate conformation is likely to correspond to the drug-bound state. The intermediate conformation is similar to the 1PBC-bound structure. Using AWH, the free energy minima reliably capture the binding configurations of 1PBC, A9C, niclosamide, and Ani9. The binding site of 1PBC, A9C, and niclosamide agrees well with the previous computational and experimental mutagenesis data ^6,19,37^. This approach is very similar to the metadynamics applied to the Nav channel family blockers ^27^. While the free energy values differ somewhat far from experimental data owing to limited convergence, the free energy landscape nonetheless provides useful information about the binding mode of the ligand in its pocket.

The power of the enhanced sampling on the structure obtained from the minima has granted a clearer picture of the niclosamide binding site. Until now, the binding site of niclosamide on the TMEM16x family has been solved only for the TMEM16F lipid scramblase ^7^, or by docking studies on the TMEM16A structure^29^. The binding site in AWH overlaps with the highly conserved site on TMEM16F. While this work suggests that niclosamide has a different site of action in TMEM16A than in TMEM16F, the sites follow a common logic. In TMEM16F, niclosamide binds at the pre-scrambling lipid headgroup binding site, while the site in TMEM16A is located at the chloride entry point.

A major challenge in drug design is subtype specificity. Ani9 is specific for TMEM16A but not for TMEM16B. TMEM16B also exhibits less conductance than TMEM16A (0.8 vs 8.3 pS, respectively) ^10,12^. Interestingly, PIP_2_ activates the TMEM16A channel but inhibits TMEM16B ^12^. Yet, 1PBC can block both subtypes ^6^. These observations led to the hypothesis that TMEM16B has a constricted pore and, thus, lower conductance. Using AWH and FAST, we show that Ani9 may exhibit dual binding sites at both the lower and outer gates of TMEM16A. As the outer gate is more constricted on TMEM16B, there is no binding site for Ani9. In addition, the absence of S592 in TMEM16B hinders the ability of the ability to bind intracellularly. Altogether, this work suggests that the conductive state of TMEM16B more closely resembles the intermediate state, or pore-blocker-bound state, of TMEM16A, given its accessibility to smaller drug molecules.

Recent years have brought major advances in the discovery of TMEM16A-specific drug interactions through physics- and AI-based screening ^37–39^, complemented by multiple recent structures of the TMEM16A channels. The power of adaptive sampling enables physics-based conformational sampling, which may allow us to access conformations unavailable via an AI-based approach, as is common for ion channels, including the Cys-loop receptors ^34^. Overall, this study highlights the power of adaptive sampling, particularly FAST, to capture both the intermediate and conductive states of TMEM16A. Endogenous ligands, such as PIP_2_, provide a powerful computational tool for stabilising the protein in a conformation which has not been captured experimentally. The AWH results emphasised that the conductive state of an ion channel is not always the druggable state, underscoring the importance of protein conformations in rational drug design for ion channels in the new paradigm of combined physics- and AI-driven drug discovery.

## Methods

### System preparation and molecular dynamics simulation

The human TMEM16A (Uniprot ID: W6JLH6) model was generated from the structure of mouse TMEM16A with Ca^2+^ bound (PDB entry: 5oyb), or with 1PBC (PDB entry: 7zk3) using Swissmodel^40^ from residues 117-135, 158-471 and 480-910. Throughout this manuscript and the Supplementary Information, residues are numbered according to the mouse TMEM16A structures (PDB 5oyb and 7zk3), which is the numbering used in the TMEM16A structural and mutagenesis literature, so that our models can be compared directly with the experimentally determined structures. Note that the model itself was built from the human TMEM16A sequence; human numbering is offset by +26 relative to this numbering for residues C-terminal to residue 481 (owing to a 26-residue insertion present in the human sequence), and is identical N-terminal to that point. Thus the outer-gate residues V543 and I640, the pore-lining Y514 and R515, and the pore lysine K645 correspond to V569, I666, Y540, R541 and K671 in human numbering. Where the residue identity differs between the two species, this is stated explicitly in the text. The structure of TMEM16B with Ca2+ bound was co-folded using the sequence from Uniprot ID Q8CFW1 with two calcium ions using AlphaFold3^41^. The proteins were converted to the MARTINI2.2 coarse-grained force field using martinize.py ^42^, embedded in a palmitoyl oleyl phosphatidylcholine (POPC) bilayer using insane.py, and equilibrated for 50 ns^43^. All simulation parameters and setups with different lipid compositions are provided in the SI Appendix. To identify the PIP2 binding site, the simulations were conducted for 10 μs. The binding site was obtained using PyLIPID, and the site with the highest occupancy and residence time was chosen ^44^. The system was then converted to all-atom using CG2AT2 with the CHARMM36m force field, under 0.15 M NaCl and TIP3P water model ^45,46^. PIP_2_ was modelled in its fully deprotonated, −4 net-charge state using the standard CHARMM36m PIP_2_ parameters. During the coarse-grained production run, the protein backbone was position-restrained in the x, y and z directions at 1000 kJ mol−1 nm−2 (define = -DPOSRES) so that the cryo-EM geometry was preserved while the elastic network equilibrated, enabling accurate CG2AT2 back- mapping to the all-atom structure. The PIP_2_ parameter is set to the CHARMM36m force field, with a charge of -4. The parameters for the pore blockers were generated using Ligand Reader and Modeller on CHARMM-GUI ^47^. All simulations were conducted with GROMACS 2024.3. In the simulations with an applied electric field, the field was applied externally along the z-axis with no oscillation, with the field strength specified in *SI Appendix Table 1*.

### Coarse-grained alchemical free-energy calculations

The binding free energy of PIP_2_ was calculated using coarse-grained alchemical free-energy calculations, following the protocol established previously^48^. Briefly, the PIP_2_ molecule bound at the TM3–TM4–TM5 site was decoupled from the protein over 16 equally spaced λ windows. Each window was simulated for 15 ns under the MARTINI2.2, with the first 2 ns being discarded, and 15 independent repeats were performed. Free energies were obtained with the multistate Bennett acceptance-ratio estimator (MBAR) as implemented in alchemlyb/pymbar. Convergence was assessed from the MBAR overlap matrix (Supplementary Fig. 2), and the reported ΔG is the mean ± standard error across the 15 repeats.

### Adaptive sampling and MSM building

The FAST algorithm was used to adaptively sample the conductive state of the TMEM16A channel. The distance between the residues lining TM3, TM4, and TM6, listed in the SI appendix, was maximised over 15 iterations and minimised over 10 iterations. Each iteration was run for 50 ns, with a frame step of 0.5 ns. The Markov-state model (MSM) was built for each iteration, yielding no more than 500 clusters. Upon building the Markov State model, the free energy landscape was clustered into three distinct macrostates after 25 simulation cycles (10 towards closing and 15 towards opening). The MSM were built with a lag time of 10 ns, 10 dimensions, 200 k-means clusters, and 3 macrostates, as determined by the CK test. Each cluster centre was then ranked with equal weight given to the rarity of the state and the degree to which the selected distance was minimised. The Gaussian width of 0.36 was used to distinguish between each state. The top 10 states were then reseeded and run until the cycle is completed. A separate tICA model was constructed for each simulation condition, as the dominant slow modes differed between conditions; states were subsequently compared across conditions using structural metrics (outer-gate distance, TM6 kink angle, and the pore-facing orientation of Y514 and R515). All MSM building and calculations were performed using the PyEMMA 2.5.7 package ^49^.

### Accelerated weighted histogram

The free energy of drug accessibility to the binding site within the TMEM16A channel was calculated using AWH^50^. Pressure was maintained at 1 bar, and temperature was maintained at 310 K. An independent AWH bias was applied for each equilibrated structure, simulating 4 walkers for 1.5 μs, sharing bias data and target distribution. Bias was applied along the z-axis, defined relative to the centre of mass of the drug and the K645 residue in the TMEM16A channel’s vestibule, to achieve convergence. The sampling interval was 3 nm above and 1 nm below K645. The system was initialised with an average free energy error of 20 kJ/mol, a diffusion coefficient of 0.0002 nm^2^/ps, and a force constant of 12800 kJ mol^−1^ nm^−2^. A harmonic potential was applied on all Cα atoms at 5.0 kJ mol^−1^ nm^−2^. A free energy landscape was built from the AWH simulations using gmx awh.

### Sequence Alignment

Two sets of sequences (TMEM16A vs TMEM16F and TMEM16A vs TMEM16B) were aligned and compared using the Needleman-Wunsch algorithm^51^, with BLOSUM62 scoring matrix ^52^.

## Acknowledgments

T.P. thanks Dr Damon Frampton and Dr Marie Lycksell for their knowledge in electrophysiology and their support throughout this project. We thank NTU High-performance computers (NTU-HPC) for their computational support. T.P. is supported by the Lee Kuan Yew Postdoctoral Fellowship.

## Data availability statement

All tpr, itp, mdp files and the features extracted from FAST adaptive sampling are available on https://zenodo.org/records/18552571.

## Author Contributions

T.P. designed and performed research, analysed data, acquired funding and wrote the manuscript. T.P. and E.H.Y. revised the final draft of the manuscript.

