## Supplementary Fig/Table for "Accessing pore-blocker bound and alternative conductive conformations of TMEM16A using PIP_2_-assisted adaptive sampling"

**This PDF file includes:**

Supporting text

SI Methods

Supplementary Figure 1-14

Tables S1

SI References

Supporting Information Text

Supplementary methods

*Model building*

The human TMEM16A (Uniprot ID: W6JLH6) model was generated from the structure of mouse TMEM16A with Ca2+ bound (PDB entry: 5oyb) (1), or with 1PBC (PDB entry: 7zk3) (2) using Swissmodel (3) from residue 117-135, 158-471 and 480-910. The structure of TMEM16B with Ca2+ bound was co-folded using the sequence from Uniprot ID Q8CFW1 with two calcium ions using AlphaFold3 (4). Residues are numbered according to the mouse TMEM16A structures (PDB 5oyb, 7zk3), as used in the TMEM16A structural literature; the model was built from the human TMEM16A sequence, for which numbering is +26 for residues C-terminal to 481.

*Coarse-grained molecular dynamics simulation*

The proteins were converted to the MARTINI2.2 coarse-grained force field using *martinize.py (1)* with the elastic network cutoff at 0.8 nm. The protein was then embedded in a palmitoyl oleyl phosphatidylcholine (POPC) bilayer, or the system with 90% POPC and 10% palmitoyl oleyl phosphatidylinositol-4,5-bisphosphate (POP2), using *insane.py.* The system was energy minimised using the steepest descent algorithm, and equilibrated for 50 ns (*2*). The system was equilibrated at 323 K with v-rescale temperature coupling (*3*), and the pressure was maintained at 1.0 bar using Parrinello-Rahman pressure coupling (*4*). To identify PIP_2_ binding site, the simulations were conducted for 10 μs. The binding site was obtained using PyLIPID, and the site with the highest occupancy and residence time was chosen (*5*).

*All-atoms molecular dynamics simulation*

The system is then converted to all-atom using CG2AT2 with the CHARMM36m force field (*6, 7*). All simulations were conducted with GROMACS 2024.3. In all simulations, the system was energy minimised using the steepest descent algorithm and equilibrated with a restraint on Cα atoms at 1000 kJ mol^−1^ nm^−2^ for 10 ns. The temperature was maintained at 310 K using a v-rescale thermostat, and 1 atm semi-isotropic pressure coupling was maintained using a c-rescale barostat (*8*). In the simulations with an applied electric field, it is applied externally along the z-axis with no oscillation. The list of all-atom simulations conducted is shown in Table 1.

*Adaptive sampling and MSM building*

The FAST algorithm was used to adaptively sample the open state of the TMEM16A channel (13). The distance between the carbon atoms on the Cα and the side chain of the residues lining (1) TM3 (Q637, I641, G644, K645 and Q649), TM4,6 (V543, L547, V548, V549, I551, D554, E555, A600, K603,); (2) TM4 (V543, L547, V548, V549, I551, D554, E555) and TM6 (V599, A600, K603); and (3) E555-L652 and E555-E654 was maximised over 15 iterations and minimised over 10 iterations. The minimum distances between the same pair of residues were used to build the MSM. Each iteration was run for 50 ns, with a frame step of 0.5 ns. The Markov-state model (MSM) was built for each iteration, with the cluster radius dynamically adjusted, yielding no more than 500 clusters. The Gaussian width of 0.36 was used to distinguish between each state. Upon building the Markov State model, the free energy landscape was clustered into three distinct macrostates after 25 simulation cycles (10 towards closing and 15 towards opening). The MSM were built with a lag time of 10 ns, 10 dimensions, 200 k-means clusters, and 3 macrostates, as determined based on implied timescale plot and the CK test. Each cluster centre was then ranked with equal weight given to the rarity of the state and the degree to which the selected distance was minimised. The top 10 states were then reseeded and run until the cycle is completed. All MSM building and calculations were performed using the PyEMMA 2.5.7 package (14).

*Accelerated weighted histogram*

The free energy of drug accessibility to the binding site within the TMEM16A channel was calculated using AWH (*9*). Pressure was maintained at 1 bar, and temperature at 310 K. An independent AWH bias was applied to each equilibrated structure, simulating 4 walkers for 1.5 μs, sharing the bias data and target distribution. Bias was applied along the z-axis, defined relative to the centre of mass of the drug and the K645 residue in the TMEM16A channel's vestibule, to achieve convergence. Sampling interval was 3 nm above and 1 nm below K645. The system was initialised with an average free energy error of 20 kJ/mol, a diffusion coefficient of 0.0002 nm^2^/ps, and a force constant of 12800 kJ mol^−1^ nm^−2^. A harmonic potential was applied on all C_α_ atoms at 5.0 kJ mol^−1^ nm^−2^. A free energy landscape was built from the AWH simulations using *gmx awh*.

Figures

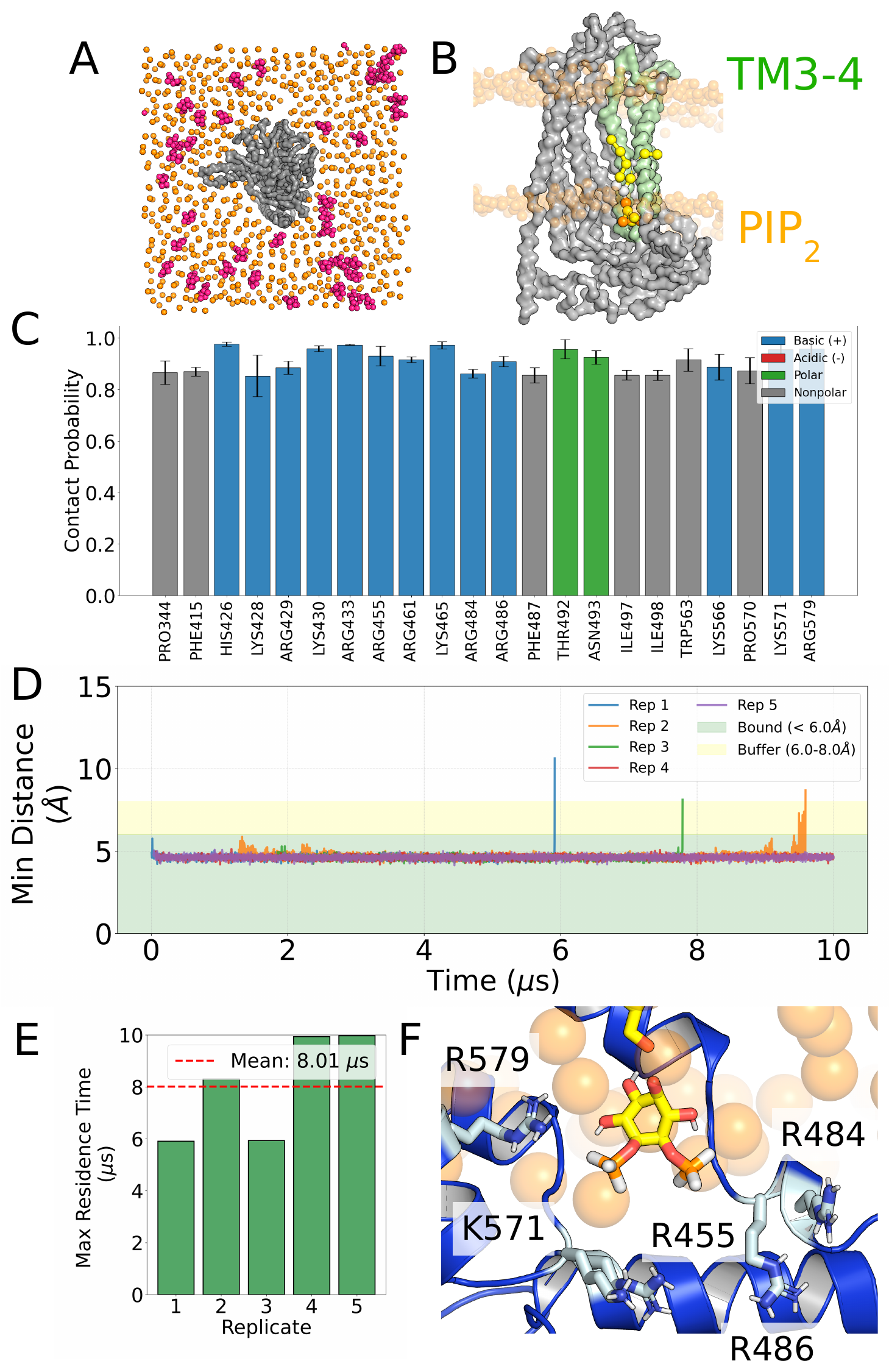

**Supplementary Fig. 1. Identification of the PIP2 binding site using coarse-grained MD simulation.**

(A) Set-up for the identification of the PIP_2_ binding site, with TMEM16A (grey surface) embedded in a phospholipid bilayer (orange spheres, P atoms only) containing 10% PIP2 in the lower leaflet (pink spheres). (B) PIP_2_ binding site on the TMEM16A channel (grey surface), highlighting the importance of TM3–TM4 (green) and PIP_2_ (yellow spheres). (C) Residues making more than 0.85 contact probability with PIP_2_ across five 10 μs simulation repeats; error bars show the standard deviation. (D) Minimum distance between the PIP_2_ headgroup and every residue of the coarse-grained protein, with each repeat shown in a different colour. Green shading marks the primary 6 Å cut-off and yellow shading the secondary 8 Å (rattling) cut-off. (E) Residence time of PIP2 within the binding site for each repeat. (F) Representative binding pose of PIP2 (yellow sticks) in the binding site (blue) after conversion back to an all-atom representation using CG2AT2.

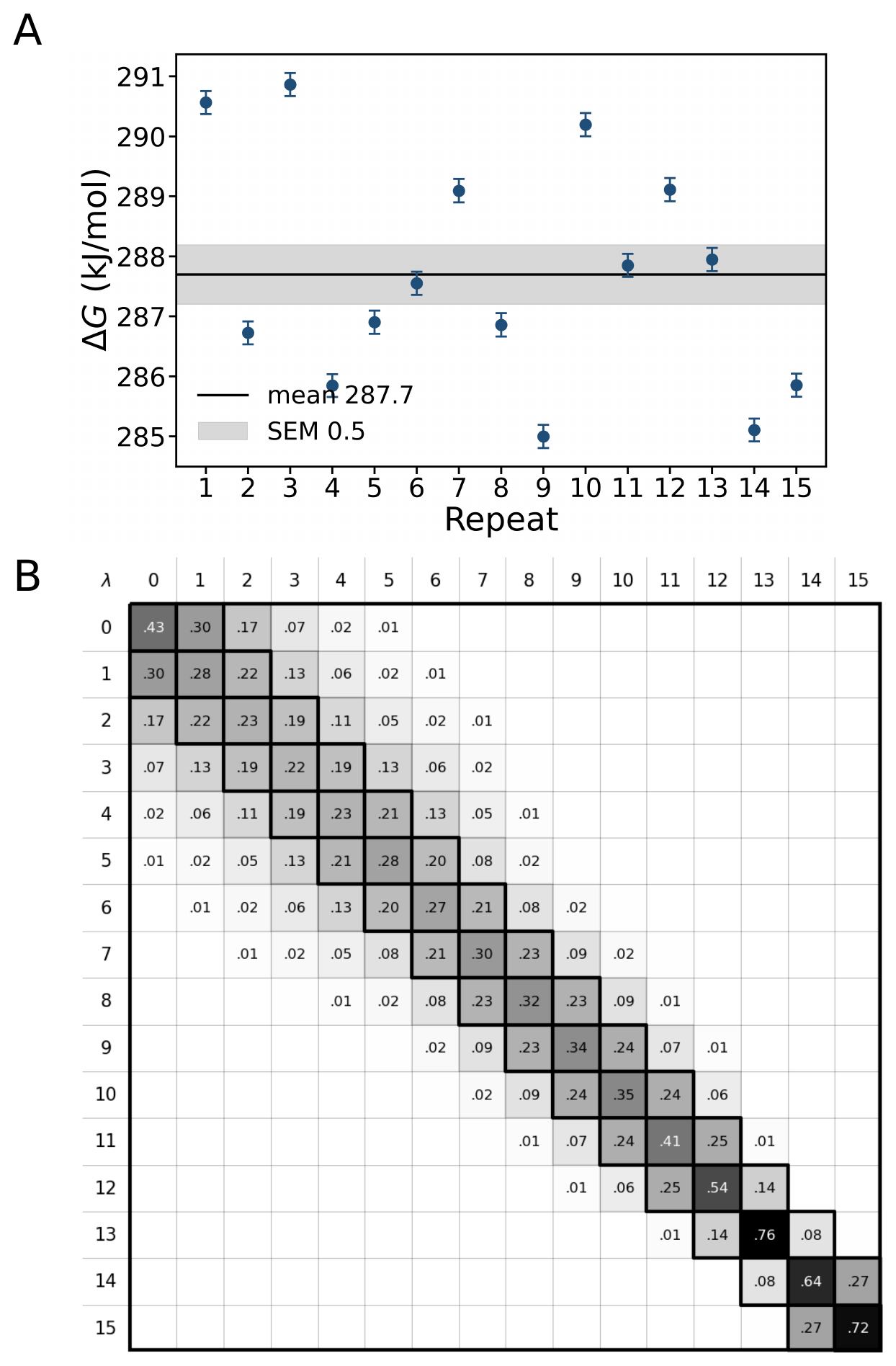

**Supplementary Fig. 2. PIP2 binding free energy from alchemical free-energy perturbation.**

(A) Coupling free energy of PIP_2_ at the TM3–TM4–TM5 site, computed by alchemical free-energy perturbation and analysed with MBAR over 15 independent repeats, each comprising 16 λ windows. Points show the per-repeat ΔG with their MBAR uncertainties; the solid line and grey band denote the mean ± SEM across repeats (287.7 ± 0.5 kJ mol^−1^). (B) MBAR overlap matrix between neighbouring λ windows (λ = 0–15); the dominant diagonal and non-zero off-diagonal elements confirm sufficient phase-space overlap for a reliable free-energy estimate.

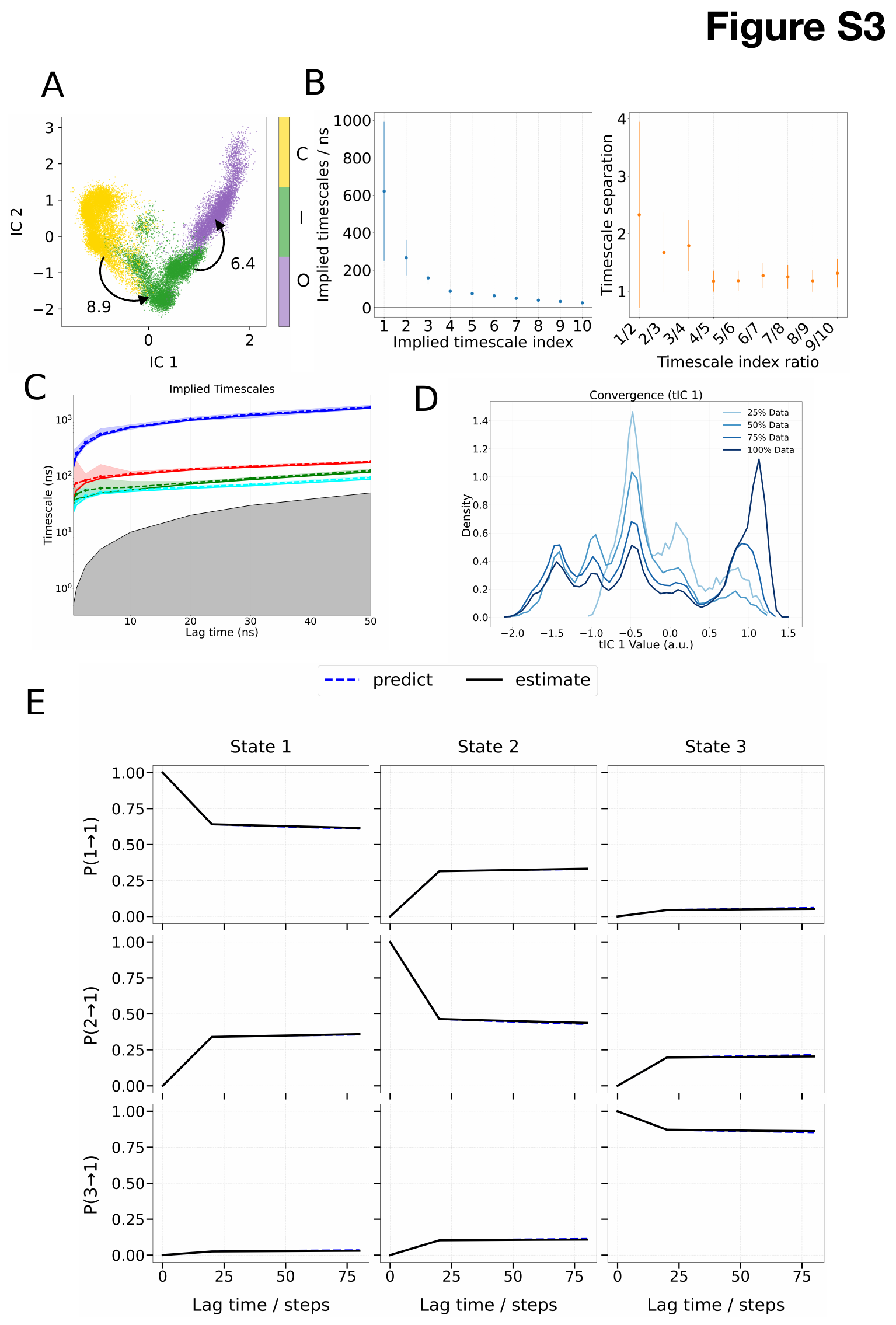

**Supplementary Fig. 3. Implied timescales and convergence of the Markov state model for TMEM16A with Ca^2+^ and PIP_2_.**

(A) Projection of the sampled conformations, coloured by PCCA macrostate: purple, open (O); green, intermediate (I); yellow, closed (C). Numbers indicate the mean first passage time in ms and each dot represents a single simulation frame. (B) Implied timescales (ns) for the top 10 collective variables (tICs); the separation shows that the slowest conformational change is captured within the first four tICs. Error bars denote 95% confidence intervals. (C) Implied timescales for the four slowest processes plotted against lag time. Solid lines represent the maximum-likelihood estimate (MLE) and dashed lines the ensemble mean from Bayesian sampling; shaded regions denote 95% confidence intervals. The curves converge at a lag time of ~10 ns, indicating the onset of Markovian behaviour. Grey shading marks the resolution limit, where timescales are faster than the lag time. (D) Sampling density along the first tIC, showing that the transition region is thoroughly sampled by at least 75% of the data. (E) Chapman–Kolmogorov test for the three-macrostate MSM, comparing estimated transition probabilities (solid) with model predictions (dashed); each step is 0.5 ns. The overlap indicates that the model has converged.

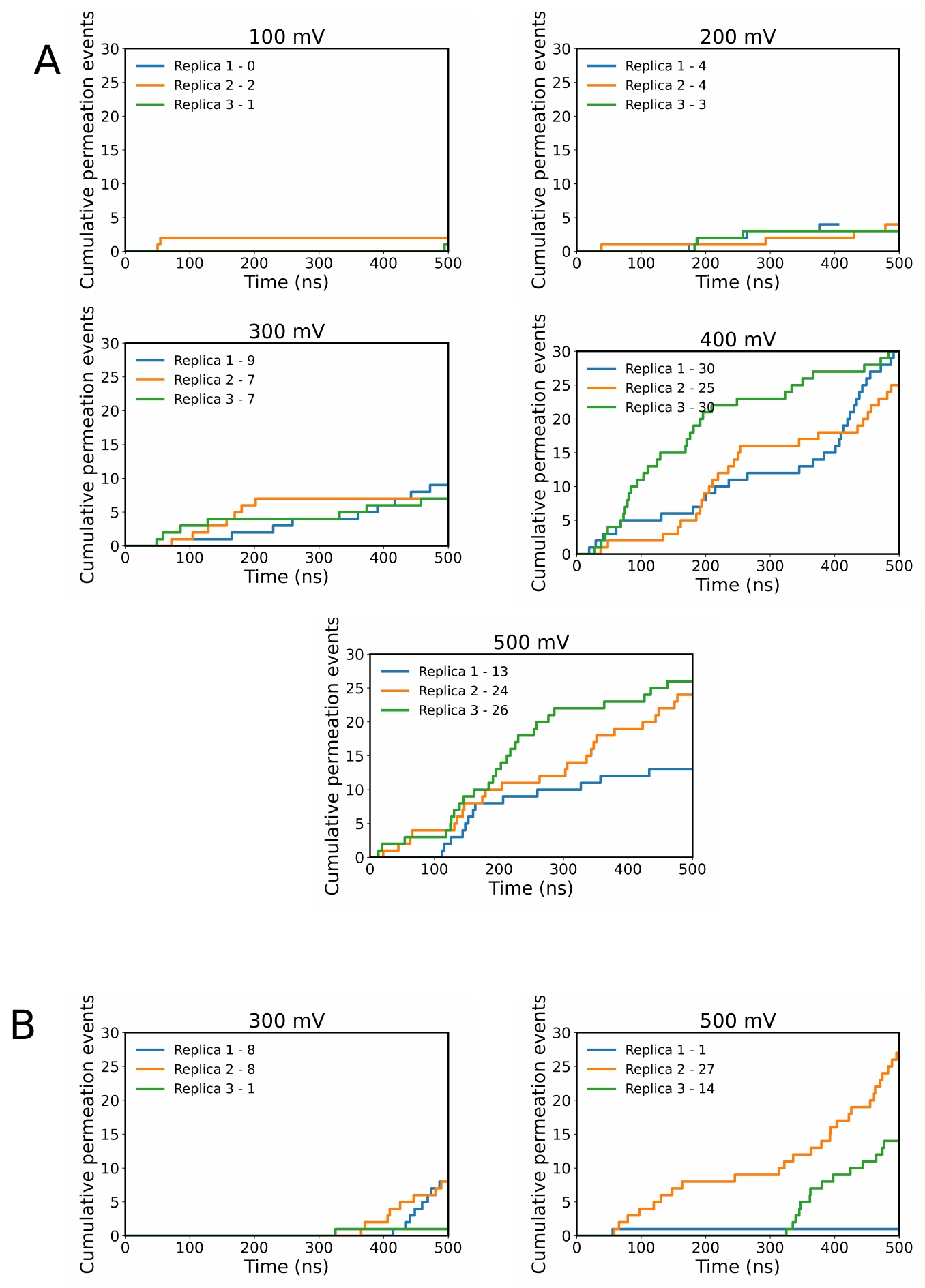

**Supplementary Fig. 4. Computational electrophysiology of the TMEM16A channel.**

(A) Cumulative chloride permeation events across the conductive (O) state of TMEM16A under applied electric fields of +100, +200, +300, +400 and +500 mV. Different colours indicate independent repeats and the total event count for each replica is given in the legend. Permeation is infrequent at +100 mV (0–2 events per replica), so the current carries a large counting uncertainty, whereas at +200–300 mV the counts rise and are consistent across the three replicas. (B) Cumulative chloride permeation across the 1PBC-bound TMEM16A structure (PDB 7zk3) under +300 and +500 mV, for comparison.

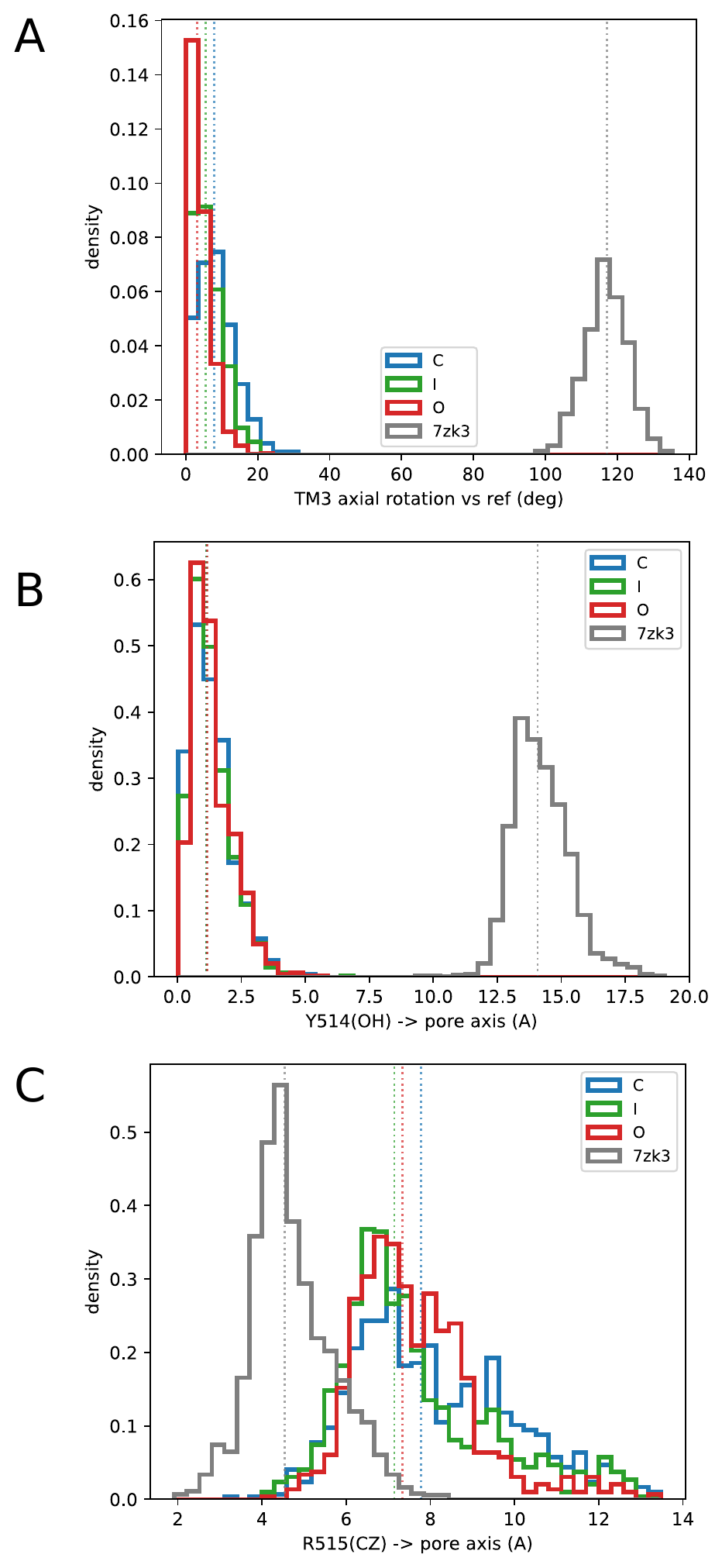

**Supplementary Fig. 5. Comparison of the pore geometry of the FAST-derived states with the 1PBC-bound structure (7zk3).**

Distributions of three intrinsic gate parameters for the closed (C, blue), intermediate (I, green) and conductive (O, red) states, compared with a reference simulation of the 1PBC-bound structure 7zk3 (grey). (Top) Axial rotation of TM3 relative to the reference; the FAST-derived states do not reproduce the large TM3 rotation seen in 7zk3. (Middle) Distance from the Y514 (Y514 in mouse) hydroxyl to the pore axis; Y514 remains oriented towards the pore in all three states, unlike in 7zk3. (Bottom) Distance from the R515 (R515 in mouse) guanidinium carbon (Cζ) to the pore axis, showing only a partial shift of R515 in the conductive and intermediate states relative to 7zk3. Dashed vertical lines indicate the distribution means.

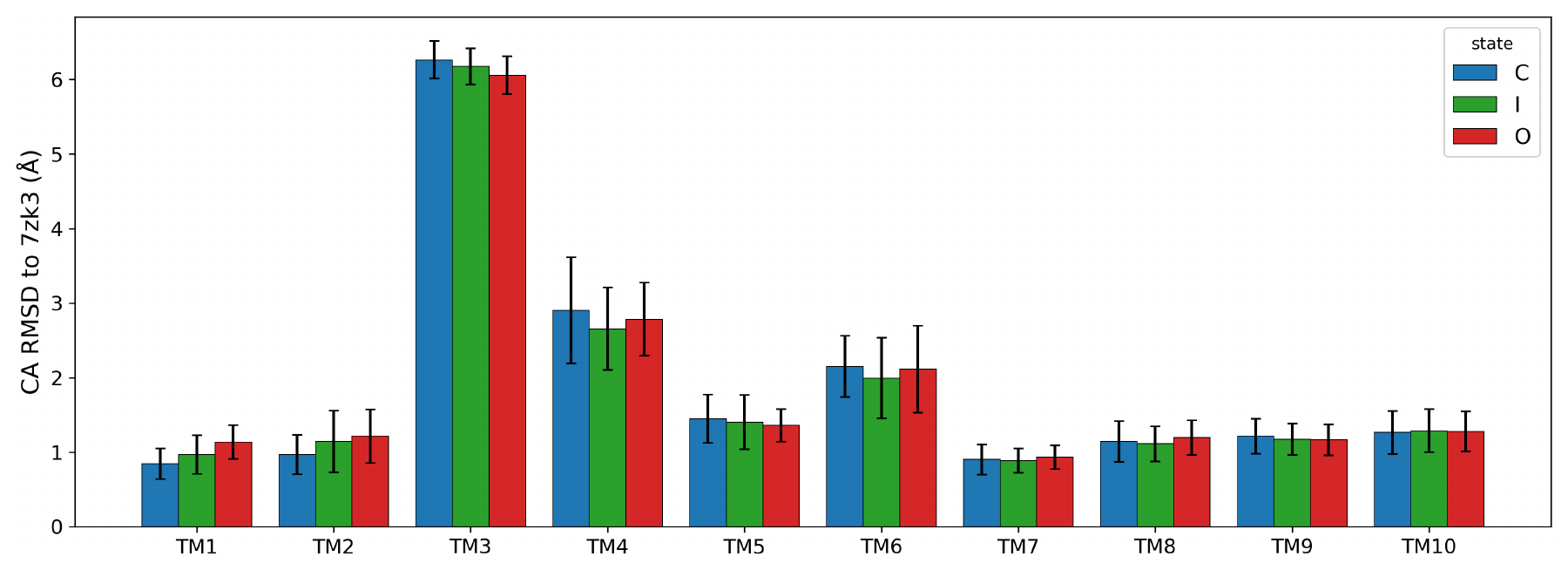

**Supplementary Fig. 6. Per-helix backbone RMSD of each state to the 1PBC-bound structure (7zk3).**

Cα RMSD of each transmembrane helix (TM1–TM10) to 7zk3 after superposition of the complete transmembrane bundle, for the closed (C, blue), intermediate (I, green) and conductive (O, red) states. Bars show the mean and error bars the standard deviation across frames. All helices lie within ~1–3 Å of 7zk3 with the exception of TM3, which deviates by ~6 Å in every state and is therefore the sole structural outlier, consistent with the absence of the axial TM3 rotation in our models.

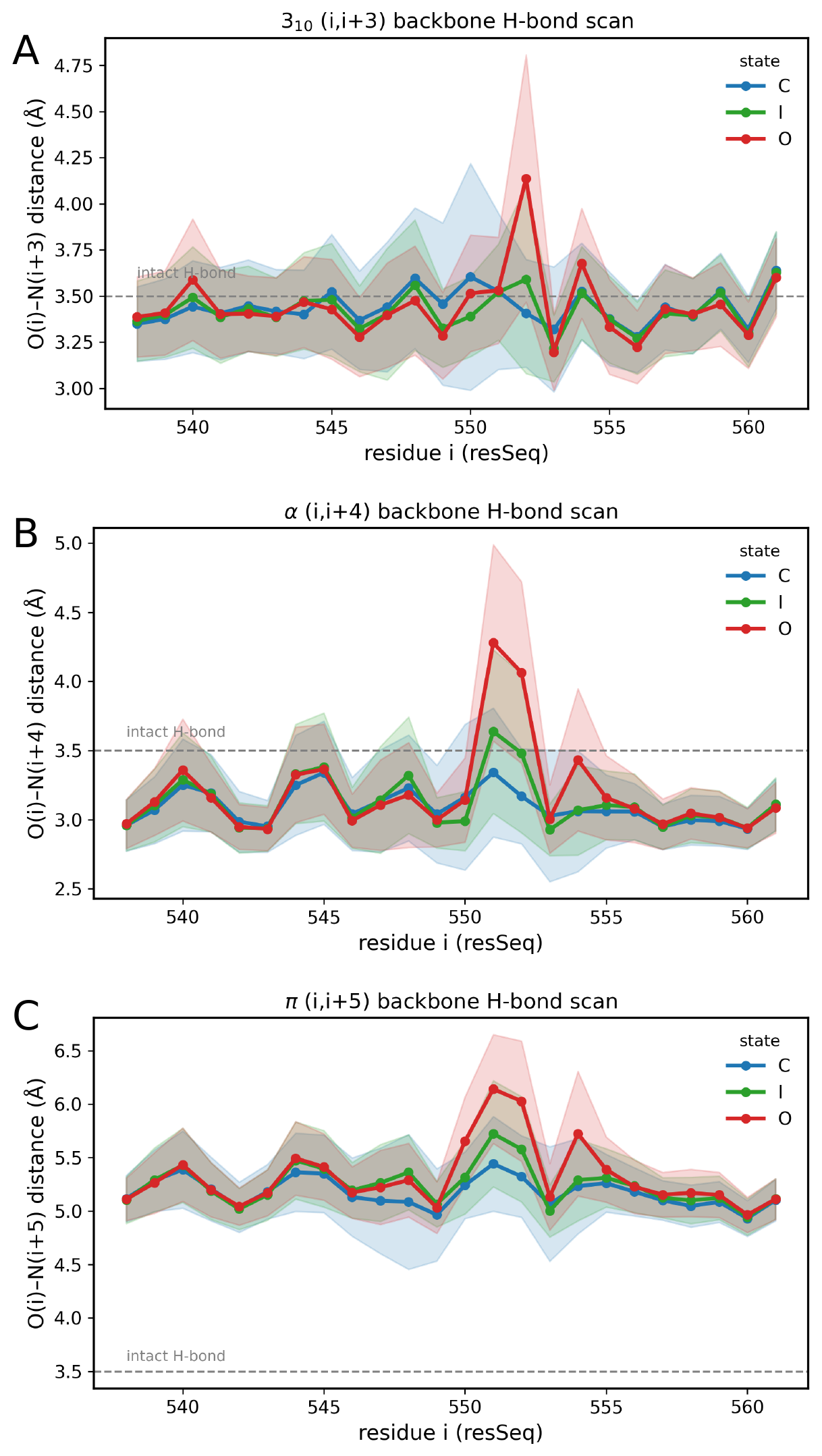

**Supplementary Fig. 7. Backbone hydrogen-bond analysis of the α-to-π transition in TM4.**

Backbone hydrogen-bond distance scans across residues 564–587 of TM4 for the closed (C, blue), intermediate (I, green) and conductive (O, red) states. (Top) 310-helical (i, i+3) O(i)–N(i+3) distances. (Middle) α-helical (i, i+4) O(i)–N(i+4) distances. (Bottom) π-helical (i, i+5) O(i)–N(i+5) distances. Lines show the mean and shaded regions the standard deviation; the dashed grey line marks the 3.5 Å intact hydrogen-bond criterion. The conductive state shows a local loss of the α-helical hydrogen bond and a corresponding gain of π-helical character around residues 550–553, consistent with an α-to-π transition in TM4.

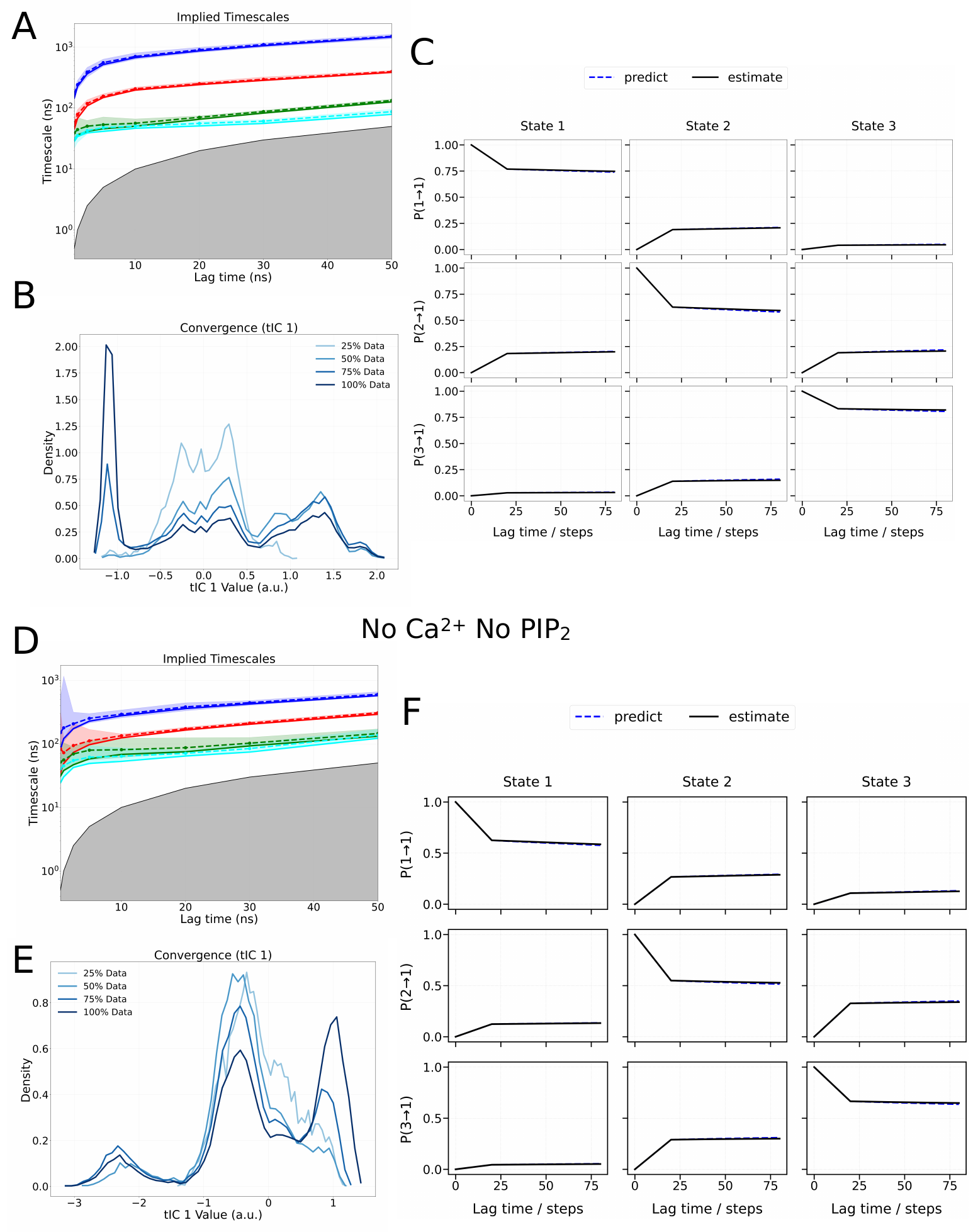

**Supplementary Fig. 8. Implied timescales and convergence of the Markov state models for TMEM16A without PIP_2_.**

Simulations in the presence of Ca2+ (A–C) and in the absence of Ca2+ (D–F). (A, D) Implied timescales for the four slowest processes (blue, red, green and cyan) plotted against lag times of 0.5, 1, 5, 10 and 25 ns; convergence at ~10 ns indicates Markovian behaviour, and grey shading marks timescales that cannot be resolved. (B, E) Sampling density along the first tIC, showing that the transition region is sampled by at least 75% of the data. (C, F) Chapman–Kolmogorov test for the three-macrostate MSM, comparing estimated transition probabilities (solid) with model predictions (dashed); the overlap indicates convergence.

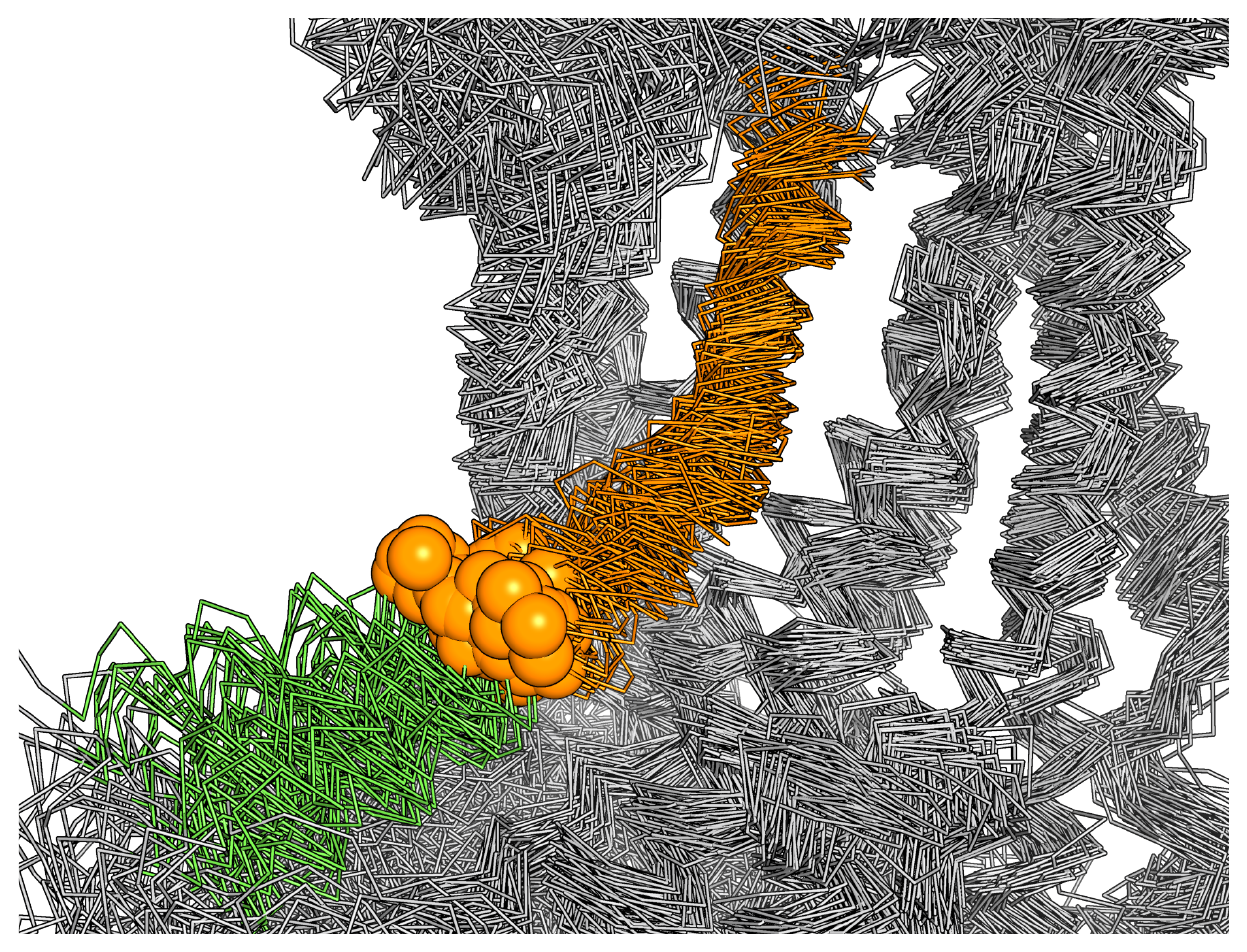

**Supplementary Fig. 9. Flexibility of the cytoplasmic end of TM6.**

Superposition of representative snapshots taken from the free-energy minima, aligned on the structurally stable membrane core (TM3, TM4, TM5, TM7 and TM8; grey). TM6 is shown in orange and the adjacent cytoplasmic helix in green, with the bound Ca2+ ions as orange spheres. Whereas the aligned core remains essentially invariant, the cytoplasmic half of TM6 samples a broad range of positions, illustrating the intrinsic flexibility of the lower part of TM6 about the G644 kink.

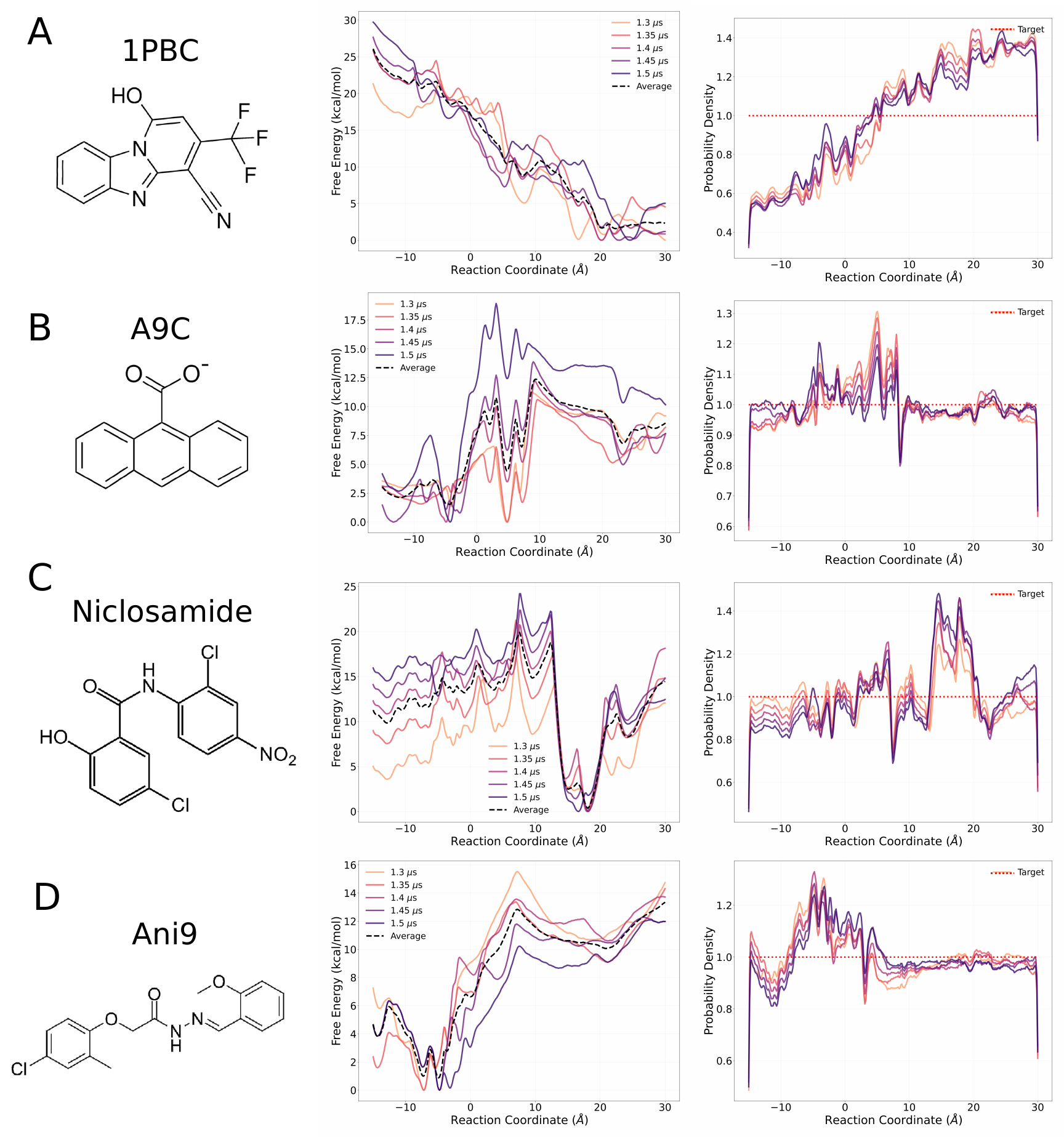

**Supplementary Fig. 10. Convergence of the accelerated weight histogram (AWH) simulations.**

Convergence of the AWH sampling along the z-axis for (A) 1PBC, (B) A9C, (C) niclosamide and (D) Ani9. (Left) Chemical structure of each drug molecule. (Middle) Free-energy landscape projected along the centre-of-mass distance between the drug and K645 along the z-axis, shown at 1.30, 1.35, 1.40, 1.45 and 1.50 μs in different colours; the dashed line is the average free-energy landscape. (Right) Coordinate distribution of the sampling at the same time points, with the target distribution shown as a dashed red line.

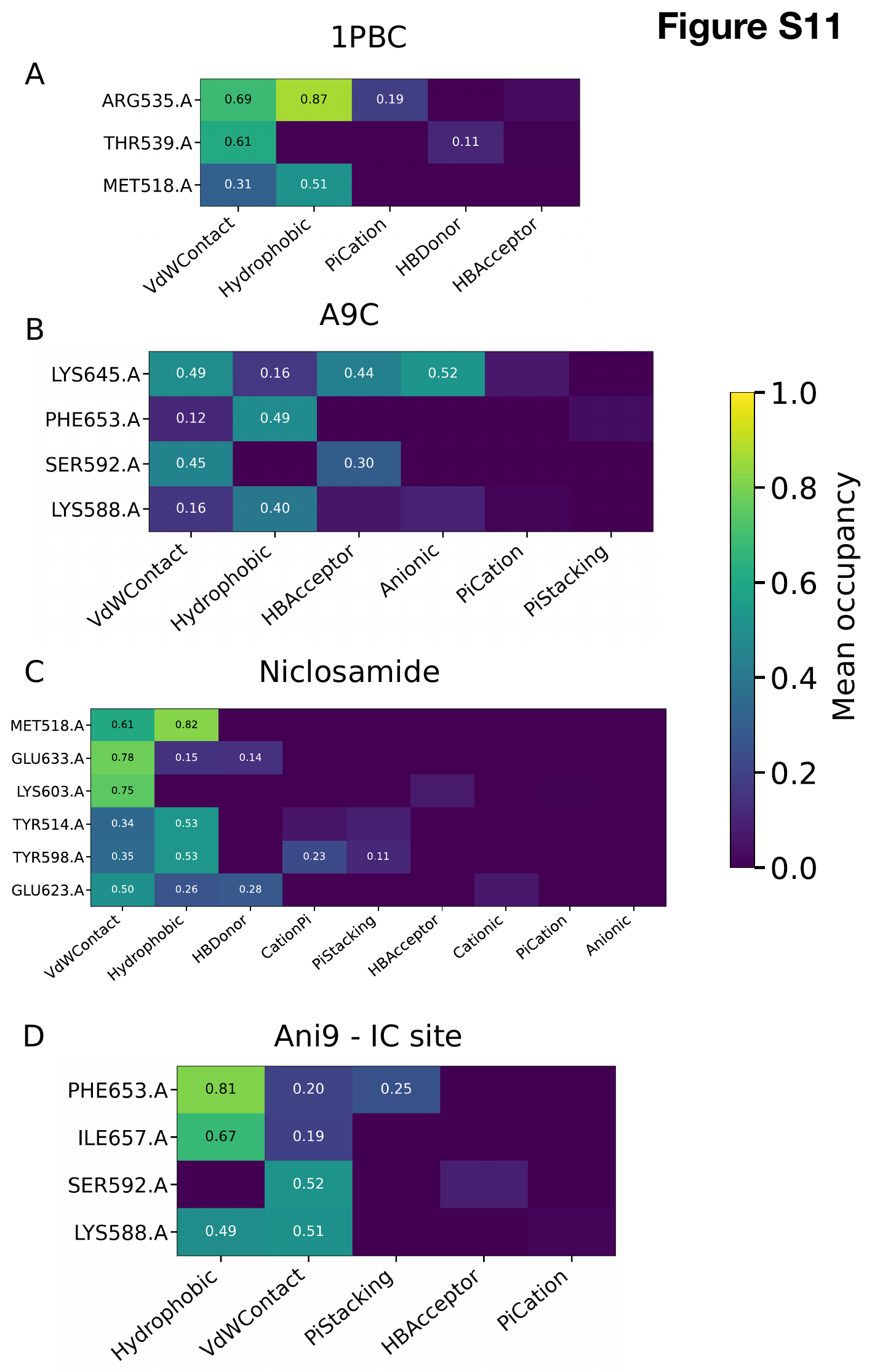

**Supplementary Fig. 11. Protein–ligand interaction fingerprints for the four pore blockers.**

Per-residue non-bonded interaction occupancies computed with ProLIF over the dominant binding-pose cluster of (A) 1PBC, (B) A9C, (C) niclosamide and (D) Ani9 at the intracellular site. Colour indicates the mean occupancy, i.e. the fraction of frames in which each interaction is present, and interactions are separated by type (hydrophobic, hydrogen bond, π-stacking, van der Waals and ionic contacts). Residues are numbered according to the TMEM16A (7zk3) sequence.

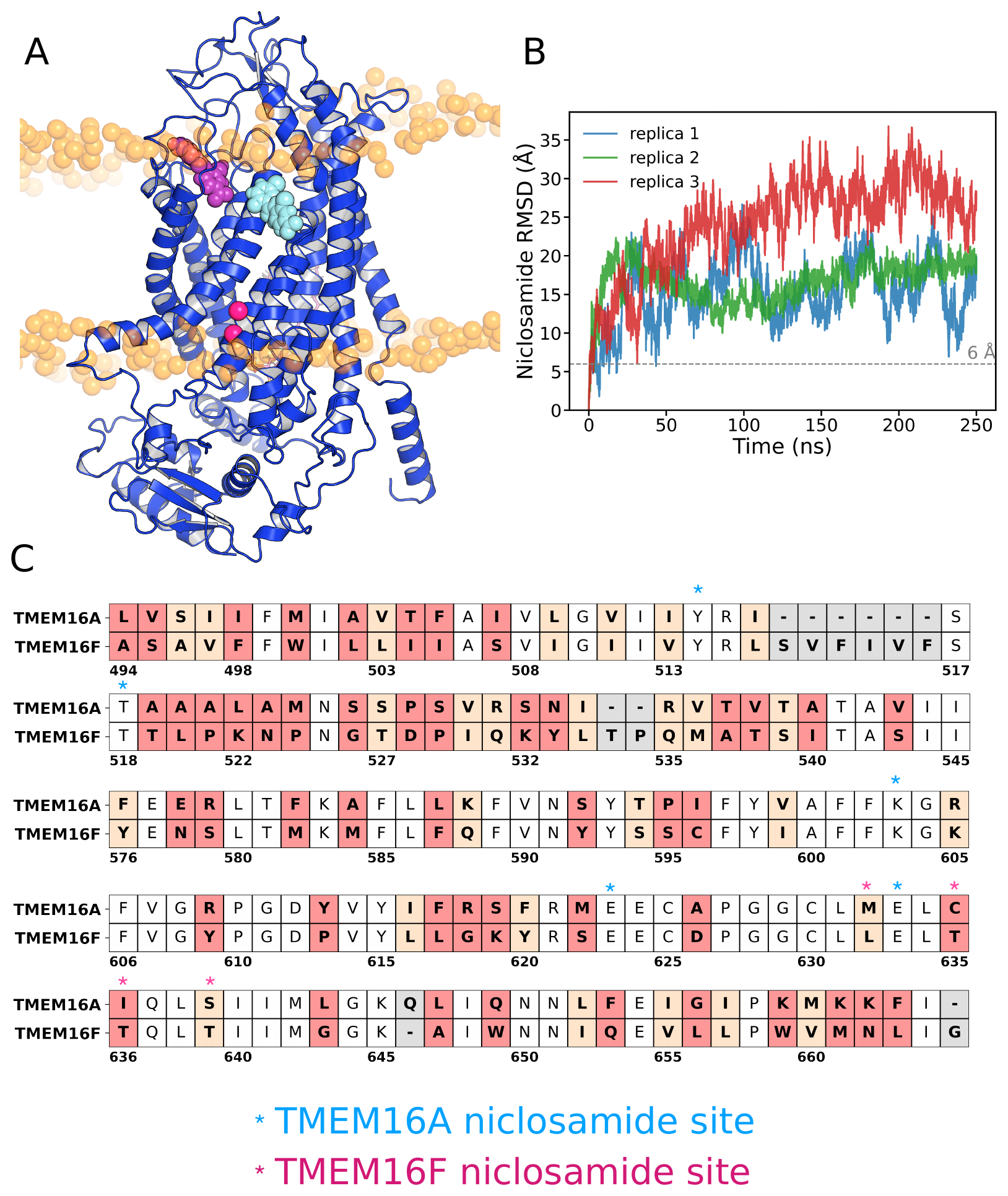

**Supplementary Fig. 12. Niclosamide binding site on TMEM16A compared with TMEM16F.**

(A) Niclosamide binding site obtained from the TMEM16F cryo-EM structure (cyan) and from the AWH simulations of TMEM16A (purple), shown on the TMEM16A channel (blue). Phosphorus atoms of the lipid phosphate headgroups are shown as orange spheres. (B) Root-mean-square deviation of niclosamide initiated from the cryo-EM TMEM16F pose in TMEM16A; the ligand leaves this shallow site in every replica, indicating that the TMEM16F pose is not stable in TMEM16A. (C) Sequence alignment of TMEM16A and TMEM16F generated with the Needleman–Wunsch algorithm and the BLOSUM62 scoring matrix. Orange marks partially conserved (medium-scoring) regions and red marks low-scoring regions; asterisks indicate the residues forming the niclosamide site in TMEM16A and in TMEM16F.

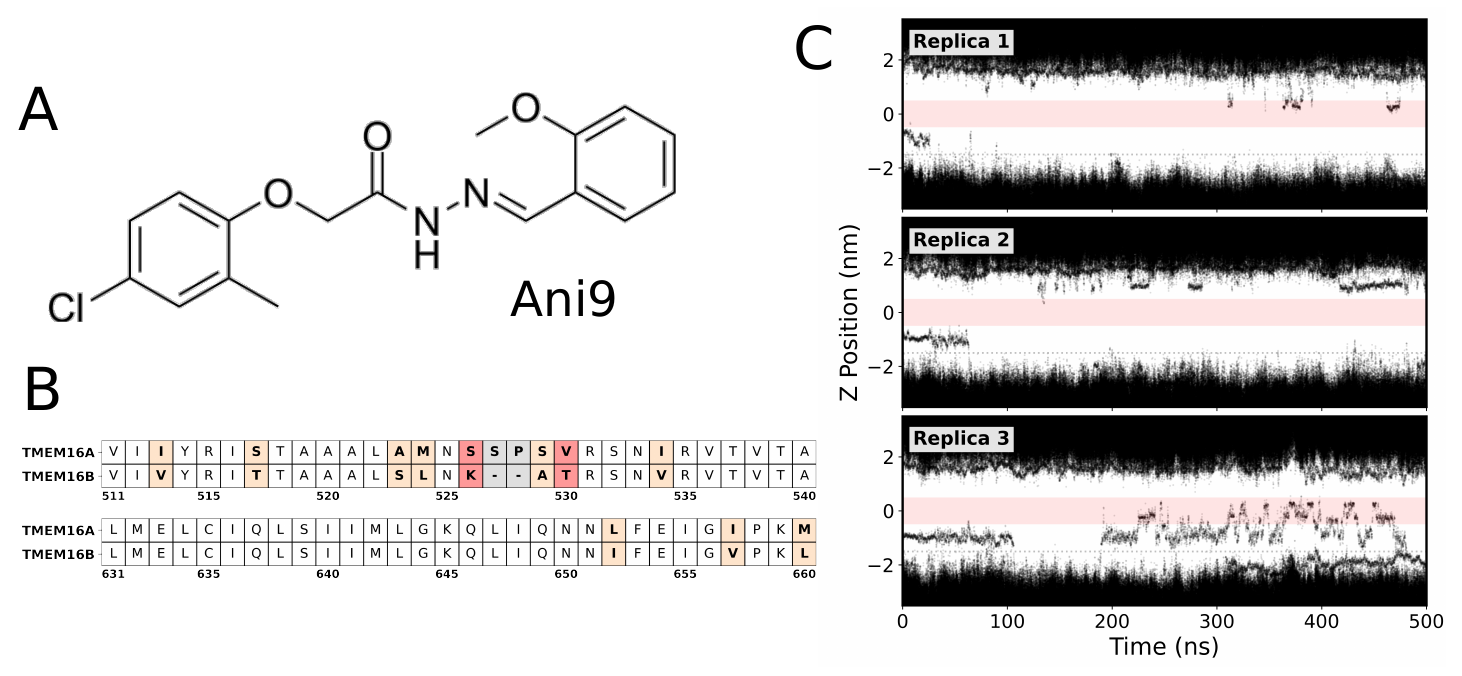

**Supplementary Fig. 13. The outer pore of TMEM16A and TMEM16B is highly conserved.**

(A) Chemical structure of Ani9. (B) Sequence alignment of TMEM16A and TMEM16B generated with the Needleman–Wunsch algorithm and the BLOSUM62 scoring matrix. Orange marks partially conserved (medium-scoring) regions and red marks low-scoring regions. (C) Position of chloride ions (black) along the pore axis across three 500 ns repeats of the conductive-state channel with Ani9 bound under a +300 mV electric field. Red shading marks the transmembrane region and position 0 corresponds to K645.

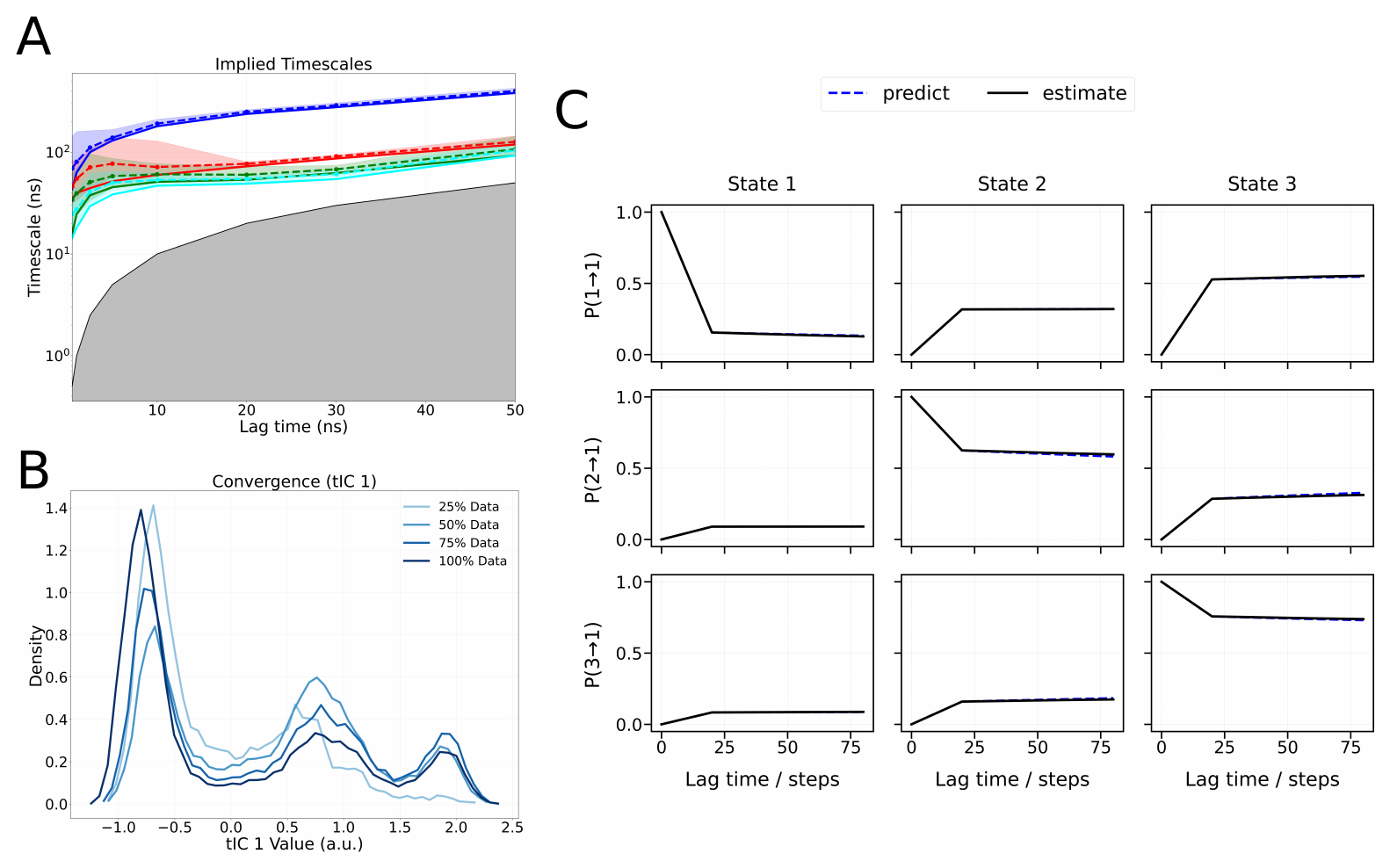

**Supplementary Fig. 14. Implied timescales and convergence of the Markov state model for TMEM16B with Ca2+.**

(A) Implied timescales for the four slowest processes (blue, red, green and cyan) plotted against lag times of 0.5, 1, 5, 10 and 25 ns. Convergence at a lag time of ~10 ns indicates Markovian behaviour; grey shading marks timescales that cannot be resolved. (B) Sampling density along the first tIC using 25%, 50%, 75% and 100% of the data, showing that the transition region is thoroughly sampled. (C) Chapman–Kolmogorov test for the three-macrostate MSM, comparing estimated transition probabilities (solid) with model predictions (dashed dark blue); the overlap indicates that the model has converged.

Tables

Table S1. All-atoms molecular dynamics simulations conducted

| **System and conditions** | **Time (ns)** | **Repeats**  **(gen x kids)** | **Aggregated time**  **(μs)** |
| --- | --- | --- | --- |
| PIP_2_ + Ca^2+^ 0 mV | 50 | 25 x 10 | 12.5 |
| Ca^2+^ no PIP_2_ 0 mV | 50 | 25 x 10 | 12.5 |
| No Ca^2+^ no PIP_2_ 0 mV | 50 | 25 x 10 | 12.5 |
| TMEM16B + Ca^2+^ 0 mV | 50 | 25 x 10 | 12.5 |
| O state 100mV | 500 | 1x3 | 1.5 |
| O state 200mV | 500 | 1x3 | 1.5 |
| O state 300mV | 500 | 1x3 | 1.5 |
| O state 400mV | 500 | 1x3 | 1.5 |
| O state 500mV | 500 | 1x3 | 1.5 |
| I state 100mV | 500 | 1x3 | 1.5 |
| I state 200mV | 500 | 1x3 | 1.5 |
| I state 300mV | 500 | 1x3 | 1.5 |
| I state 400mV | 500 | 1x3 | 1.5 |
| I state 500mV | 500 | 1x3 | 1.5 |
| 7zk3 300mV | 500 | 1x3 | 1.5 |
| 7zk3 500mV | 500 | 1x3 | 1.5 |
| O state + 1PBC 0 mV | 250 | 1x3 | 0.75 |
| I state AWH + 1PBC 0 mV | 1500 | 1x4 | 6 |
| I state AWH + A9C 0 mV | 1500 | 1x4 | 6 |
| O state + Niclosamide 500mV | 500 | 1x5 | 2.5 |
| O state + Niclosamide 0 mV | 250 | 1x5 | 1.25 |
| I state + Niclosamide 0 mV | 250 | 1x5 | 1.25 |
| I state AWH + Niclosamide 0 mV | 1500 | 1x4 | 6 |
| O state + Ani9 300mV | 500 | 1x3 | 1.5 |
| I state AWH + Ani9 0 mV | 1500 | 1x4 | 6 |
|  |  | **Total** | **99.25** |
